# Asymmetric neural dynamics of visuospatial attention in autism spectrum disorder

**DOI:** 10.64898/2026.06.04.730213

**Authors:** Megan Darrell, Theo Vanneau, Chloe Brittenham, John J. Foxe, Sophie Molholm

## Abstract

**Background:** Selective attention enables the prioritization of behaviorally relevant information in complex sensory environments. Despite substantial evidence for altered attention in autism spectrum disorder (ASD), the neurophysiological mechanisms underlying these differences remain poorly understood.

**Methods:** Here, we integrate high-density electroencephalography (EEG), pupillometry, and behavioral measures collected during a cued covert visuospatial selective attention task to characterize mechanisms of spatial attention in children and adolescents with ASD (n = 18; 13.4 ± 3.0 YO), and how they differ from age- and IQ- matched individuals with typical-development (TD) (n = 21; 14.7 ± 3.8 YO).

**Results:** Both groups demonstrated high target detection accuracy and comparable response times, with no significant between-group differences in behavioral performance. Furthermore, neurophysiological measures demonstrated that during leftward attention, both TD and ASD participants exhibited canonical attentional processes, including lateralized anticipatory parieto-occipital alpha modulation and enhanced P1 sensory responses to attended stimuli. Additionally, across both groups, trial-level analyses revealed that decreased anticipatory alpha power and increased P1 amplitude contralateral to the attended hemifield were associated with faster reaction times. In contrast, there were notable group differences in the neural dynamics supporting rightward spatial attention. TD participants showed early sensory gain (P1 modulation) without alpha-band modulation, whereas ASD participants exhibited modulation of posterior alpha power without effective sensory gain. Interestingly, for rightward attention, only P1 amplitude predicted reaction time, and this was the case for both groups. Resting-state alpha dynamics did not differ between groups, indicating that the attended hemifield differences reflect task-dependent differences in attentional control rather than baseline oscillatory differences.

**Limitations:** Limitations include modest sample size and restriction to autistic individuals with relatively low support needs, which may limit the generalizability of these findings to the broader autism spectrum.

**Conclusions:** The similarity of leftward attention mechanisms across groups, which includes intact recruitment of anticipatory alpha modulation, argues against a global disruption of basic visuo-spatial attentional function in autistic individuals with low support needs. However, group differences emerged specifically during rightward attention, where ASD participants showed a more uniform pattern of oscillatory modulation, warranting further investigation. Collectively, these findings provide novel insight into the neural architecture of visuospatial attention in ASD, revealing how preparatory oscillatory activity shapes early sensory responses and behavior during selective attention.

## Introduction

In complex sensory environments, the brain is confronted with far more information than it has the capacity to process (Desimone and Duncan 1995). Selective attention provides a mechanism for navigating this challenge, dynamically selecting which signals are prioritized while simultaneously suppressing others (James 1890). When this process functions atypically, differences in how information is prioritized can influence the filtering of irrelevant input, thereby shaping how individuals experience and engage with their environment.

Mounting evidence suggests that these filtering mechanisms are altered in autism spectrum disorder (ASD), where attentional differences are highly prevalent. Estimates suggest that 40–70% of individuals with ASD—compared to approximately 10.5% of U.S. children overall (Danielson, Claussen et al. 2024)—meet criteria for attention-deficit/hyperactivity disorder (ADHD) (Joshi, Faraone et al. 2017, Lyall, Croen et al. 2017, Brookman-Frazee, Stadnick et al. 2018), underscoring the central role of attentional atypicality in this population. Empirical findings, which are largely derived from behavioral studies, often characterize attention in ASD as more broadly distributed or hypo-focused, reflecting inefficient filtering of irrelevant stimuli (Remington, Swettenham et al. 2009, Remington, Swettenham et al. 2012, Murphy, Foxe et al. 2014, Keehn, Nair et al. 2016). Difficulty filtering distracting stimuli likely contributes to core clinical symptoms in ASD, including sensory overarousal (Liss, Saulnier et al. 2006, Brinkert and Remington 2020) and difficulties in social interactions (Falck-Ytter, Kleberg et al. 2023).

While behavioral findings highlight the prevalence of atypical attention in ASD, the neural mechanisms underlying these differences remain poorly understood. Electroencephalography (EEG) provides a powerful tool to probe these mechanisms by capturing temporally resolved neural dynamics that index attentional control. Converging evidence suggests that distributed attentional control is coordinated through neural oscillations, which reflect synchronized modulation of neural activity (Petersen, Robinson et al. 1987, Zhou, Schafer et al. 2016), particularly alpha-band activity (∼7-13 Hz), which is often characterized as the dominant intrinsic rhythm of the human brain ((Berger 1929); see (E Basar 1996, Basar, Schurmann et al. 1997) for review) and is also widely thought to play a central role in attentional control (Foxe and Snyder 2011, Hanslmayr, Gross et al. 2011, Bonnefond and Jensen 2025). Increases in alpha power over posterior cortex—usually measured in the anticipatory interval between an attentional cue and the imperative stimulus—have been associated with suppression of unattended spatial locations (Worden, Foxe et al. 2000, Kelly, Gomez-Ramirez et al. 2009), distracting visual input during inter-sensory attention tasks (Foxe, Simpson et al. 1998, Fu, Foxe et al. 2001, Gomez-Ramirez, Kelly et al. 2011), and task-irrelevant visual features such as color or motion (Snyder and Foxe 2010)). Importantly, these alpha dynamics also scale with behavioral performance on the primary task (Thut, Nietzel et al. 2006, Kelly, Gomez-Ramirez et al. 2009, Murphy, Foxe et al. 2014). According to this framework, fluctuations in alpha-band amplitude mediate cortical suppression: periods of elevated alpha power reflect inhibitory states associated with sensory suppression, whereas reduced alpha power corresponds to a release of inhibition and increased cortical excitability (Klimesch, Sauseng et al. 2007, Klimesch, Sauseng et al. 2007, Romei, Gross et al. 2010, Hanslmayr, Gross et al. 2011, Wilson and Foxe 2020, Han, Brincat et al. 2026).

During perceptually “optimal” states, when posterior alpha power is low and local cortical inhibition is reduced, the visual system is thought to enter a state of sensory sampling that facilitates target detection. Under these conditions, increased cortical excitability should manifest as greater sensory gain at the neural level, indexed by the stimulus-evoked response. Indeed, electrophysiological and neuroimaging studies consistently show stronger neural responses to attended compared with unattended stimuli (Moran and Desimone 1985, Hillyard, Vogel et al. 1998). These attentional enhancements typically emerge 80–200 ms after stimulus onset (P1 and N1 components) (Luck, Heinze et al. 1990, Heinze, Mangun et al. 1994, Wijers, Lange et al. 1997, Woldorff, Fox et al. 1997, Kelly, Gomez-Ramirez et al. 2008) and are thought to reflect enhanced perceptual processing under conditions of attentional demand (Awh, Anllo-Vento et al. 2000).

Given the role of these oscillatory and sensory gain mechanisms in typical attentional control, it remains unclear whether these processes function similarly in ASD. Previous EEG and functional magnetic resonance imaging (fMRI) studies offer preliminary evidence of atypical neural mechanisms supporting selective attention and distractor suppression in ASD (Townsend and Courchesne 1994, Teder-Salejarvi, Pierce et al. 2005, Kawakubo, Kasai et al. 2007, Ohta, Yamada et al. 2012, Keehn, Westerfield et al. 2017). Here, we consider the possibility that difficulties filtering distracting sensory information in ASD may reflect atypical alpha-band dynamics, consistent with reports of reduced anticipatory alpha suppression during inter-sensory selective attention tasks (Frey, Molholm et al. 2013, Murphy, Foxe et al. 2014) and diminished alpha modulation associated with inefficient filtering of irrelevant stimuli (Keehn, Nair et al. 2016, Keehn, Westerfield et al. 2017). Furthermore, resting alpha-band power has also been shown to be significantly altered in ASD (see (Neo, Foti et al. 2023) for review), although previous studies report conflicting findings regarding its directionality (higher (Ogawa, Sugiyama et al. 1982, Sutton, Burnette et al. 2005) or lower (Chan, Sze et al. 2007, Murias, Webb et al. 2007, Keehn, Westerfield et al. 2017) in ASD). Accordingly, task-related differences in alpha modulation may stem from underlying differences in resting-state alpha activity present prior to task engagement.

Despite mounting evidence for altered alpha activity, the functional integrity of alpha suppression mechanisms for attentional control in ASD is not well understood. Therefore, to address these gaps, we conducted a comprehensive investigation of selective visuospatial attention in autistic and typically-developing (TD) children and adolescents, integrating high-density EEG with pupillometry and clinical assessments of attention and autism-related symptomatology. Within this framework, we aimed to elucidate the neurophysiological mechanisms supporting visuospatial attention in ASD and determine how these mechanisms relate to clinical phenotypes. Moreover, despite social impairment being a core diagnostic feature of ASD—and extensive evidence of reduced social orienting and atypical face processing across behavioral studies ((Osterling and Dawson 1994), see (Chita-Tegmark 2016) for review), neuroimaging and electrophysiological investigations (Dawson, Carver et al. 2002, Schultz, Grelotti et al. 2003, Webb, Dawson et al. 2006, van Kooten, Palmen et al. 2008, Webb, Jones et al. 2011, Vanneau, Brittenham et al. 2025)—there is a relative paucity of research investigating neural mechanisms of attention in social contexts in ASD. We therefore included both social (faces) and non-social (houses) stimuli to determine whether the neural dynamics supporting spatial attention in ASD are modulated by social context. Together, this multimodal approach provides a comprehensive characterization of the neural dynamics supporting spatial attention in autistic and TD youth.

## Methods

### Participants

Data from 27 TD and 26 ASD participants were initially collected. Of these, four TD and one ASD participants were excluded prior to preprocessing due to eye-tracking equipment incompatibilities (e.g., calibration failure due to glasses). Three ASD participants were excluded due to poor behavioral performance (<50% accuracy in any block or incomplete task completion).

Following EEG and eye-tracking preprocessing, one TD and two ASD participants were excluded due to excessive EEG noise (>15% bad channels or >50% bad epochs), and an additional one TD and two ASD participants were excluded due to excessive saccades (>50% of trials exceeding 2° outside the central fixation region of interest; see *Eye-Tracking Preprocessing*). The final sample included 21 TD and 18 ASD participants (**Table 1**).

**Table 1.**
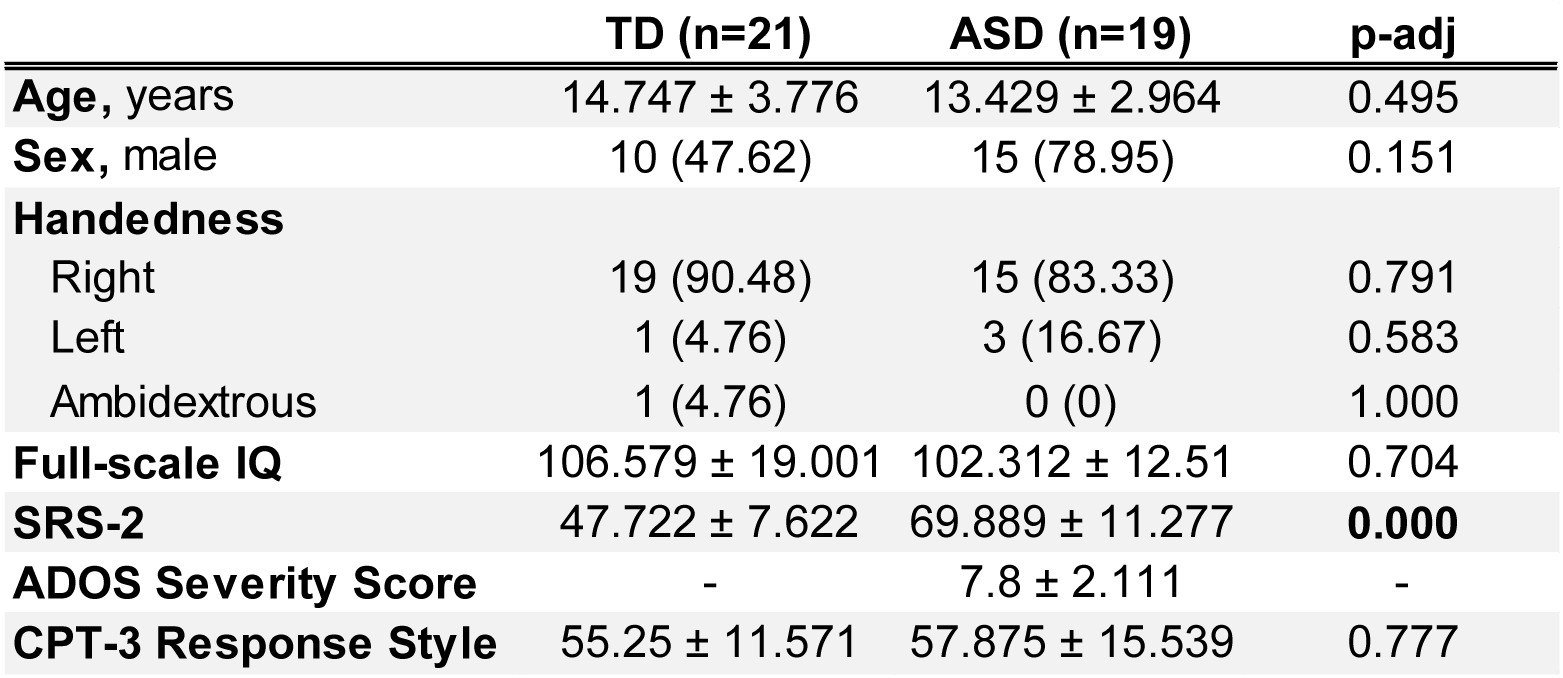
Participant demographics. Continuous variables: mean ± standard deviation. Categorical variables: n (%). Missing clinical data: IQ (TD: n=19, ASD: n=16); SRS-2 (TD: n=16; ASD: n=18); CPT-3 (TD: n=20; ASD: n=16); Handedness (ASD: n=20); ADOS-2 (ASD: n=16) *IQ – intelligence quotient; SRS-2 – Social Responsiveness Scale, 2^nd^ edition; CPT-3 – Conners Continuous Performance Task, 3^rd^ edition*

|  | TD (n=21) | ASD (n=19) | p-adj |
| --- | --- | --- | --- |
| **Age, years** | 14.747 ± 3.776 | 13.429 ± 2.964 | 0.495 |
| **Sex, male** | 10 (47.62) | 15 (78.95) | 0.151 |
| **Handedness** |  |  |  |
| Right | 19 (90.48) | 15 (83.33) | 0.791 |
| Left | 1 (4.76) | 3 (16.67) | 0.583 |
| Ambidextrous | 1 (4.76) | 0 (0) | 1.000 |
| **Full-scale IQ** | 106.579 ± 19.001 | 102.312 ± 12.51 | 0.704 |
| **SRS-2** | 47.722 ± 7.622 | 69.889 ± 11.277 | **0.000** |
| **ADOS Severity Score** | - | 7.8 ± 2.111 | - |
| **CPT-3 Response Style** | 55.25 ± 11.571 | 57.875 ± 15.539 | 0.777 |

Participants were recruited without regard to sex, race, or ethnicity; Intelligence quotients for verbal (V-IQ), and full-scale (FS-IQ) intelligence were assessed for all participants, using the Wechsler Abbreviated Scales of Intelligence (WASI-II; (Weschler 2011)) and Wechsler Intelligence Scale for Children (WISC-V; (Weschler 2014)).

The Social Responsiveness Score, 2^nd^ Edition (SRS-2; (Constantino, Davis et al. 2003, J.N. 2012)) questionnaire was collected from all participants to obtain a continuous measure of autistic traits. The SRS-2 yields age- and sex-normed T-scores (M = 50, SD = 10), with higher scores indicating higher levels of autistic traits. T-scores ≤59 are considered within the typical range, 60–65 indicates mild impairment, 66–75 moderate impairment, and ≥76 severe impairment consistent with clinically significant symptomatology.

Continuous measures of attention were obtained from Conners’ Continuous Performance Test, 3^rd^ edition (CPT-3; (Conners 2008)), which is a computerized test to evaluate inattentiveness, impulsivity, sustained attention and vigilance in individuals >8 years old. As measured by the CPT-3, the Response Style Index is a signal detection statistic that reflects an individual’s natural response tendency in tasks involving a speed–accuracy trade-off. T-scores are age-normed (M = 50, SD = 10). A conservative response style (T ≥ 60) reflects an emphasis on accuracy over speed (i.e., a more cautious approach), whereas a liberal response style (T ≤ 40) reflects an emphasis on speed over accuracy (i.e., a more impulsive approach). T-scores between 41–59 indicate a balanced response style, reflecting sensitivity to both speed and accuracy.

To be included in the TD group, participants had to have no history of neurological, developmental, or psychiatric disorders, have no first-degree relatives with a diagnosis of ASD, and be in age-appropriate grade at school. To be included in the ASD group, participants had to meet diagnostic criteria for ASD on the basis of the following measures: *1*) autism diagnostic observation schedule 2 (ADOS-2) (Lord, Rutter et al. 1994); *2*) diagnostic criteria for autistic disorder from the *Diagnostic and Statistical Manual of Mental Disorders* (DSM-5); *3*) clinical impression of a licensed clinician with extensive experience in diagnosis and evaluation of children with ASD. Exclusionary criteria for all groups included: (1) known genetic syndrome (including syndromic cases of ASD), (2) history of or currently taking medication for seizures in the prior 2-years, (3) exclusionary physical limitations (e.g. significant vision deficits), (4) were born prematurely (<35 weeks) and/or experienced prenatal/perinatal complications, or 5) FS-IQ <80.

All participants provided informed consent or assent in accordance with age-appropriate procedures approved by the Institutional Review Board of the Albert Einstein College of Medicine. For participants under 18 years of age, written parental or guardian consent was obtained in addition to participant assent. Participants 18 years or older provided their own written informed consent Participants received modest recompense for their participation.

### Procedure

Participants completed the task while seated in a dimly lit, electrically shielded, sound-attenuated booth (International Acoustics Company, Bronx, NY). Head position and gaze were stabilized with a chin rest, and visual stimuli were presented on an LCD monitor located 79 cm from the participant.

Continuous EEG was recorded using a 64-channel BioSemi ActiveTwo system (BioSemi, Amsterdam, The Netherlands) arranged according to the international 10–20 electrode layout. Signals were digitized at 512 Hz and acquired using BioSemi’s active electrode configuration with an anti-aliasing filter (−3 dB at 3.6 kHz). During acquisition, the system used the Common Mode Sense (CMS) active electrode together with the Driven Right Leg (DRL) passive electrode to form a feedback loop that stabilizes the reference potential and suppresses common-mode noise. Unlike conventional reference electrodes, this BioSemi referencing scheme dynamically compensates for voltage differences between the scalp and amplifier ground, effectively establishing a near-zero reference potential.

The experimental paradigm was programmed in Python using PsychoPy. Event markers corresponding to stimulus onset and button responses were transmitted to the acquisition computer as analog triggers via Lab Streaming Layer (LSL) (Kothe, Shirazi et al. 2024). Eye position and pupil diameter were simultaneously recorded with an EyeLink 1000 eye-tracking system (SR Research Ltd., Mississauga, Ontario, Canada) at a sampling rate of 500 Hz.

### Task

#### Attention Task

We adapted a traditional spatial attention paradigm that has been previously applied in children of this age (Banerjee, Snyder et al. 2011), which consists of a simple Stimulus 1 (S1) – Stimulus 2 (S2) design. Each trial consisted of an instructional cue (S1), an intervening blank preparatory period (cue-S2 window), followed by a task-relevant second stimulus (S2).

Each trial began with a 1,000 ms fixation window, followed by a centrally presented arrow cue (S1; 1.4° diameter), displayed in the center of the screen, indicating the hemifield to which participants should covertly attend while maintaining central fixation (see **Fig. 1a**). The cue–S2 stimulus onset asynchrony (SOA) was fixed at 1,120 ms. Following the cue interval, S2 stimuli were presented for 160 ms. These consisted of grayscale social (face from NimStim database; (Tottenham, Tanaka et al. 2009)) and non-social (house) images (9.15° × 6.62°) positioned 5.5° to the left or right of fixation. The face and house stimuli were matched for luminance, brightness, and contrast and were presented on a gray background. Including both social and non-social images allowed assessment of whether attentional sensory gain differed as a function of stimulus social relevance.

**Figure 1.**
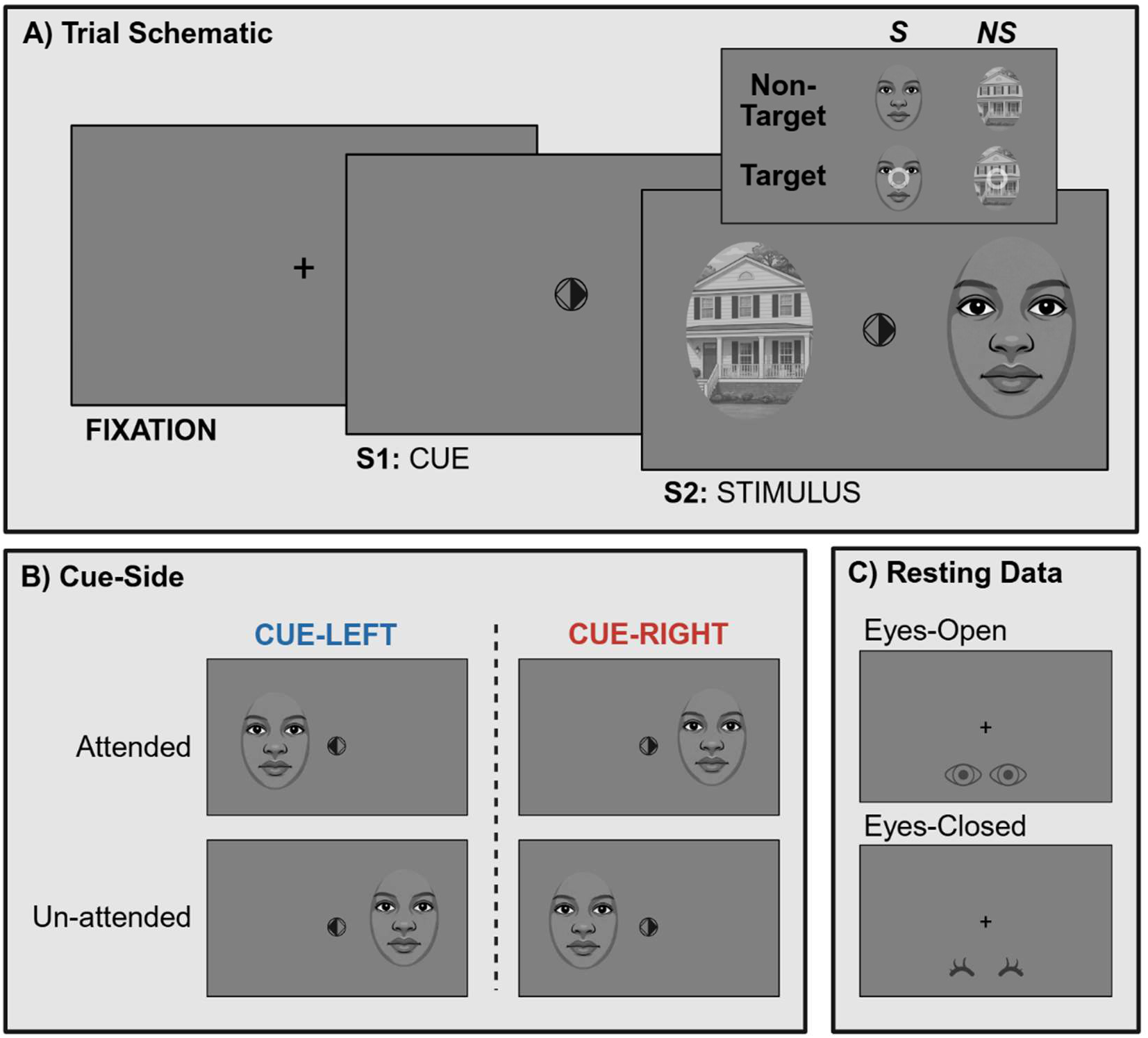
Experimental design. **a)** Trial schematic. Each trial began with a fixation period (1,000 ms), followed by a brief spatial cue (S1) indicating which side to attend to. After an anticipatory interval (1,120 ms), stimuli were presented (S2; 160 ms), consisting of social (S; faces) and/or non-social (NS; houses) images. Participants then responded via-keyboard press based on the presence of a white target ring on the attended side (response interval: 1,000 ms). This schematic is for illustrative purposes only and is not to scale. Face stimuli were originally real face images (NimStim database; (Tottenham, Tanaka et al. 2009)); for display purposes here, we show stylized comic representations to avoid identifiable likenesses. **b) Cue-side.** Attention was directed to either the left (cue-left) or right (cue-right) visual hemifield via a central arrow cue. Attended refers to stimuli presented in the cued hemifield (left for cue-left; right for cue-right), whereas unattended stimuli appeared in the opposite hemifield. Only unilateral stimulus presentations are illustrated here; however, bilateral trials were also included, in which attended and unattended stimuli were presented simultaneously. **c) Resting data.** Resting-state EEG was collected both before and after the task to characterize participants’ baseline oscillatory activity. Each resting session consisted of 2 minutes of eyes-open and 2 minutes of eyes-closed recording.

Participants were instructed to covertly deploy their attention to the stimulus in the cued hemifield and respond via keyboard press when a white target ring (inner diameter 0.90°, outer diameter 0.94°) appeared over the S2 image. Targets could occur on either side, both sides, or neither side; however, participants were instructed to respond only to targets presented on the attended side and to ignore stimuli in the uncued hemifield. Participants were asked to respond as quickly and accurately as possible, withholding responses when no target was presented on the attended stimulus. The probability of a valid target (i.e., on the attended stimulus) was 33%, and target brightness was determined individually using the Up-Down Transformed Rule (UDTR) staircase procedure described in Supplementary Methods: Equating Task Difficulty Across Participants.

The inter-trial interval (S2–Cue/response period) was fixed at 1,000 ms, during which the fixation cross remained visible. The task was performed under three stimulus contexts: non-social (houses), social (faces), and intermixed presentations of both stimulus types. The first two blocks were “pure” contexts (social or non-social; order randomized across participants). Within pure blocks, S2 stimuli were presented either unilaterally on the attended side, unilaterally on the unattended side, or bilaterally. Unilateral presentations occurred with a combined probability of 67% (attended and unattended equally likely), while bilateral presentations occurred with a probability of 33%. These conditions allowed evaluation of how expectations regarding stimulus category influence attentional modulation.

The pure blocks were followed by an inter-mixed condition in which face and house S2 stimuli appeared with equal probability on attended and unattended sides. In these blocks, S2 could occur under four presentation types for both stimulus categories, yielding eight possible combinations: unilateral attended (S, NS), unilateral unattended (S, NS), bilateral identical (S/S, NS/NS), or bilateral mixed (S/NS, NS/S). Unilateral and bilateral presentations occurred with equal probability (50% each), with attended and unattended unilateral trials equally likely. Because social and non-social stimuli were unpredictably intermixed within a block, this context minimized top-down expectations and allowed assessment of potential bottom-up attentional biases driven by stimulus category.

Each participant completed 40 blocks (9 pure social, 9 pure non-social, 22 inter-mixed) of 55 trials each, resulting in the collection of ∼167 trials per stimulus condition. Data collection for this paradigm took ∼4 hours with breaks, including a longer one in which lunch was provided.

#### Resting State

Resting-state EEG was collected before and after the task to characterize participants’ intrinsic oscillatory activity (e.g., baseline alpha power) and to examine potential changes following sustained task engagement. Each session consisted of two 2-minute blocks: eyes open and eyes closed. During the eyes-open block, participants fixated on a central marker displayed on a uniform gray screen to maintain stable gaze and minimize movement-related artifacts. For the eyes-closed block, participants were instructed to close their eyes while remaining still and relaxed. The same protocol was repeated after task completion. Because eyes-open and eyes-closed conditions provide complementary information about alpha reactivity, both were included in statistical analyses of neural dynamics at rest. Fatigue-related changes in oscillatory dynamics between pre- and post-task recordings will be examined in a separate study.

Three TD and three ASD participants were excluded from the resting-state analysis due to poor data quality (>20% bad channels) in either the before or after block, yielding a final resting-state sample of *n* = 34 (TD: *n* = 18; ASD *n* = 16).

#### Pre-Processing

EEG preprocessing procedures were identical to those previously described in Darrell et al. (2026) for the TD cohort and are additionally detailed in the Supplementary Methods (Darrell, Vanneau et al. 2026).

### Analytic approach

#### Overview

Analyses were organized around two temporal epochs: (i) the preparatory cue–S2 interval (−1000 to 0 ms), used to examine anticipatory oscillatory dynamics, and (ii) the stimulus-evoked period following S2 onset (0–1000 ms). Within the post-stimulus (S2) window, EEG and pupil responses were quantified to assess attentional modulation of early sensory gain. During the preparatory interval, analyses focused on parieto-occipital alpha-band dynamics implicated in spatially selective attention. In addition, given evidence that frontal theta rhythms coordinate large-scale attentional control, we tested whether pre-stimulus theta phase is associated with posterior oscillatory activity in the anticipatory interval and stimulus-evoked sensory responses and whether it predicts behavioral performance. All analyses additionally included group as a factor to examine differences between participants with and without ASD.

#### Behavior

Behavioral performance was quantified using reaction time (RT), hit rate, miss rate, false alarm rate, detection sensitivity (d′), and inverse efficiency score (IES). Mean RT was calculated from correct responses (hits) on valid-target (target present on attended side) trials. Hit rate was defined as the proportion of correct responses on valid-target trials. False alarm rate was calculated as the proportion of responses made on target-absent (no target present on attended side) trials. Detection sensitivity (d′) was computed as the difference between the z-transformed hit rate and false alarm rate. Inverse efficiency score (IES) was calculated as mean RT divided by hit rate, providing a composite measure of speed–accuracy trade-off, with higher values reflecting slower and/or less accurate performance

### Pupillometry: Evoked Response

#### Baseline Correction

Eye-tracking epochs were baseline-corrected using a pre-stimulus baseline window (−200 to −50 ms prior to S2 onset). Analyses focused exclusively on non-target trials to eliminate target- and motor-related confounds of the pupil response.

#### Extraction of Stimulus-Evoked Pupil Response

Peak amplitude of the stimulus-evoked pupil response was calculated as the maximum positive deflection within the S2 window (0 – 900 ms). Peak latency was extracted as the time point corresponding to this maximum value. Analysis of the pupil response were conducted only for unilateral stimulus configurations to assess attention effects without distractor confounds during bilateral stimulus presentation. To maximize statistical power, analyses were on the stimulus-evoked pupil response collapsed across pure and intermixed blocks. This decision was supported by cluster-based permutation testing of the pure–intermixed difference wave against zero, which revealed no significant effects in either TD or ASD groups (**Supplementary Fig. 1A**). Pure versus intermixed effects, which were designed to index social versus non-social distractor processing during bilateral stimulus presentations, will be evaluated in future work.

### EEG: Evoked Response

#### Baseline Correction

Event-related potentials (ERPs) were analyzed from epoched EEG data time-locked to S2 onset. Epochs were extracted from −200 to 500 ms relative to stimulus onset and baseline-corrected using a pre-stimulus baseline window (−200 to −50 ms prior to S2 onset). Analyses focused on non-target trials to isolate attentional modulation of sensory-evoked activity whilst eliminating target and motor specific responses.

#### Extraction of ERP Components

ERP analyses focused on the P1 component as an index of early sensory gain in extrastriate visual cortex. Mean P1 amplitude was computed as the average voltage within a 110–150 ms window centered on the grand-average P1 peak. Peak amplitude and latency were estimated within a broader 100–160 ms window, with peak amplitude defined as the maximum positive deflection and latency as the corresponding time point. Analyses were performed separately for unilateral and bilateral stimulus configurations.

The P1 was quantified over bilateral parieto-occipital electrode clusters defined a priori based on canonical visual ERP topographies where the P1 is maximally expressed (Luck, Heinze et al. 1990, Heinze, Mangun et al. 1994, Wijers, Lange et al. 1997, Woldorff, Fox et al. 1997, Foxe and Simpson 2002). Left hemisphere electrodes included O1, PO3, and PO7, and right hemisphere electrodes included O2, PO4, and PO8.

Analyses evaluating the evoked response to bilateral stimuli are not reported here, as bilateral trials were designed to specifically evaluate social/non-social attentional effects of distractor stimuli and are beyond the scope of the present manuscript. Evoked responses to unilateral trials were analyzed using a contralateral framework to capture maximal visual responses: stimuli presented in the right visual field were analyzed over left occipital electrodes, whereas left visual field stimuli were analyzed over right occipital electrodes. Additionally, ERP analyses were collapsed across pure and intermixed blocks to increase statistical power, as cluster-based permutation testing of the pure–intermixed difference wave against zero revealed no significant effects in either TD or ASD groups (**Supplementary Fig. 1B**).

### EEG: Time-Frequency Analysis of Oscillatory Activity

Single-trial oscillatory power and phase were derived from complex time–frequency representations computed using Morlet wavelets (MNE-Python).

#### Anticipatory Alpha Power Extraction

To quantify anticipatory alpha activity, Morlet-based time–frequency power estimates were extracted in the alpha band (7–13-Hz) for each trial, channel, and time point. Alpha power was baseline-normalized using an MNE-style log-ratio transformation:

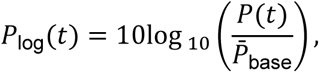

where *P*^-^_base_ denotes the per-trial mean power within a pre-stimulus baseline window (−200 to −50 ms relative to cue onset).

Attentional modulation of alpha power is typically quantified within the cue–S2 anticipatory interval. In the present study, this window was defined as −800 to −200 ms prior to S2 onset, which reflected the empirically strongest phase of preparatory modulation. Alpha power was averaged across time within this window.

Anticipatory alpha power was extracted from parieto-occipital (PO) electrodes, defined a priori based on bilateral channel groups commonly used to index visual attention–related alpha modulation (Foxe, Simpson et al. 1998, Fu, Foxe et al. 2001, Gomez-Ramirez, Higgins et al. 2007), and confirmed by the observed topographies in the current data. Specifically, the left PO ROI comprised *P3, P5, P7, PO3, PO7, O1*, and the right PO ROI comprised *P4, P6, P8, PO4, PO8, O2*.

To quantify spatially selective alpha modulation, parieto-occipital (PO) activity was analyzed within a contralateral–ipsilateral framework relative to cue direction. Trials were first separated by cue side (cue-left vs cue-right). For cue-left trials, the contralateral-cue ROI (corresponding to the attended hemifield) comprised right PO electrodes, while the ipsilateral-cue ROI (corresponding to the unattended hemifield) comprised left PO electrodes. This mapping was reversed for cue-right trials.

#### Resting-State Spectral Analysis

To determine whether task-related group differences in alpha might reflect baseline spectral differences, resting-state alpha power was extracted from the same bilateral parieto-occipital ROIs described above.

To avoid edge artifacts introduced by time–frequency convolution, analyses were restricted to the central 100-second interval of each 120-second resting block. Resting spectra were parameterized using the *specparam* (FOOOF, Fitting Oscillations & One-Over-F; (Donoghue, Haller et al. 2020)) algorithm to dissociate periodic and aperiodic components of the power spectrum. Spectral fits yielded estimates of the aperiodic exponent and periodic alpha-band peak activity. This parameterization allowed assessment of whether any hemispheric differences in alpha amplitude were attributable to oscillatory peak characteristics or to shifts in the underlying aperiodic background activity.

Resting state measures were quantified separately for hemisphere (left vs right), recording period (pre-task vs post-task), and eye condition (eyes-open vs eyes-closed), allowing direct comparison of baseline hemispheric spectral profiles independent of spatial attention demands.

### Statistical Testing

#### Per-Participant Linear Mixed-Effects Modeling (LMM): Behavioral and Neural Effects

Statistical analysis of behavioral and neural measures per-participant was conducted using a linear mixed-effects model (LMM; (Bates D 2015)) in R version 4.4.0 using the *lme4* package. Follow-up pairwise comparisons of the LMM were obtained by estimated marginal means (*emmeans*; (Lenth 2019)) to compute contrasts, applying a Tukey correction. All primary outcome measures were analyzed using LMM with subject-specific random intercepts and Age as a covariate; fixed-effect structures for each analysis are detailed in **Table 2**.

**Table 2.** LMM specifications for behavioral, pupillometric, neural, and spectral analyses. The table summarizes dependent variables and corresponding fixed-effect predictors for each primary analysis. All models included age as a covariate and subject as a random intercept (1 | Subject). Abbreviations: TD, typically-developing; ASD, autism spectrum disorder; RT, reaction time; d′, detection sensitivity; IES, inverse efficiency score; S, social; NS, non-social; IM, inter-mixed; L/R, left/right; ROI, region of interest; RS, resting state.

| Results | Measure | Predictors |
| --- | --- | --- |
| Supplementary Table 1, Fig. 2 | **Behavior** (RT, hit rate, false alarm rate, d', IES) | Group (TD/ASD) x S2 Type (unilateral/bilateral) x Block (S/NS/IM) x Cue Side (L/R) + Age + (1 \| Subject) |
| Supplementary Table 2, Fig. 3 | **Stimulus-evoked pupil response** | Group (TD/ASD) x Attention (attended/unattended) x Condition (S/NS) x Cue Side (L/R) + Age + (1 \| Subject) |
| Supplementary Table 3, Fig. 4 | **Stimulus-evoked neural response** | Group (TD/ASD) x Attention (attended/unattended) x Condition (S/NS) x Cue Side (L/R) + Age + (1 \| Subject) |
| Supplementary Table 4, Fig. 6 | **Anticipatory neuro-oscillatory power** (alpha, beta) | Group (TD/ASD) x Attention (unattended/attended ROI) x Block (S/NS/IM) x Cue Side (L/R) + Age + (1 \| Subject) |
| Fig. 7 | **Resting alpha** (power, aperiodic exponent) | Group (TD/ASD) x Hemisphere (left/right parieto-occipital ROI) x Condition (eyes-open/eyes-closed) x Resting-State (RS)-Block (before-task/after-task) + Age + (1 \| Subject) |

#### Behavioral and Data Quality Comparisons

Group differences in target ring contrast (face and house stimuli) after UDTR and in the number of retained epochs following EEG and eye-tracking preprocessing were assessed using two-sample tests. Normality was evaluated separately within each group using the Shapiro–Wilk test (Shapiro 1965). When both groups met normality assumptions (p > 0.05), group comparisons were performed using Welch’s two-sample t-tests (unequal variances); otherwise, Mann–Whitney U tests (two-sided) were used.

Target ring contrast (expressed as translucency; 1.00 = fully opaque, 0.00 = absent) after UDTR did not differ between TD and ASD groups for either face stimuli (TD: 0.356 ± 0.173; ASD: 0.334 ± 0.198; p > 0.05) or house stimuli (TD: 0.360 ± 0.217; ASD: 0.355 ± 0.229; p > 0.05). One TD and one ASD participant were excluded from this analysis because the data file containing the target contrast values was not successfully saved. Similarly, the number of retained epochs after EEG and eye-tracking preprocessing did not differ between TD (1494.2 ± 332.6) and ASD participants (1349.9 ± 441.2; p > 0.05).

#### Group Differences in Demographic and Clinical Characteristics

Group differences between ASD and TD participants in demographic and clinical variables were tested in R (v4.4.0) using two-sample procedures with α = 0.05. For continuous variables (age, full-scale IQ, SRS-2, CPT-3 response style), normality was evaluated using the Shapiro–Wilk test (Shapiro 1965). When normality was not met (p < 0.05), groups were compared using a Mann–Whitney U test (Wilcoxon rank-sum, two-sided). When normality was met, homogeneity of variance was assessed using Levene’s test (Levene 1960); variables with unequal variance (p < 0.05) were compared using Welch’s two-sample t-test, and variables with equal variance were compared using Student’s two-sample t-test (two-sided). For categorical variables (biological sex, handedness), group differences were evaluated using Pearson’s chi-square test when expected cell counts were sufficient; otherwise, Fisher’s exact test was used. Multiple comparisons across tested variables were controlled using the Benjamini–Hochberg false discovery rate procedure (Yoav Benjamini 1995).

#### Correlation analyses

Associations between neural measures and behavioral or clinical variables were assessed using correlation analyses, conducted separately within each group to avoid spurious associations driven by between-group differences. Prior to testing, variables were evaluated for normality using the Shapiro–Wilk test (Shapiro 1965). When both variables satisfied normality assumptions (p > 0.05), Pearson correlations were used; otherwise, Spearman rank correlations were applied. Correlations were calculated using pairwise complete observations.

## Results

### Behavior

As described in **Supplementary Table 1**, LMM analyses revealed consistent effects of stimulus type (unilateral versus bilateral) across behavioral measures. Hit rate showed a significant main effect (F = 4.93, p = 0.027), with higher hit rates for unilateral relative to bilateral S2 presentation (unilateral − bilateral: B = 0.024, p = 0.027). False-alarm rate also demonstrated a robust effect of stimulus type (F = 53.06, p < 0.001), reflecting fewer false alarms in unilateral compared to bilateral trials (unilateral – bilateral: B = −0.018, p < 0.001). No significant effects of stimulus type were observed for RT, inverse efficiency score (IES), or d′.

Block condition also significantly influenced behavioral performance. We identified a main effect of block type on hit rate (F = 4.64, p = 0.01), which indicated reduced hit rates in the inter-mixed relative to non-social condition (inter-mixed − non-social: B = −0.04, p = 0.01), while the social vs. non-social contrast did not reach significance (social − non-social: B = −0.029, p = 0.077). IES also demonstrated a significant main effect of block type (F = 3.07, p = 0.048). Participants exhibited higher IES in the social relative to non-social condition (social − non-social: B = 0.136, p = 0.041), indicating reduced efficiency in the social blocks; the inter-mixed vs. social contrast was not significant (inter-mixed – social: B = −0.091, p = 0.246). No significant effects of block type were observed for RT or d′.

Age was associated with reduced false-alarm rate (B = −0.002, p = 0.029) and increased detection sensitivity (B = 0.10, p = 0.001). No significant age effects were observed for hit rate, RT, or IES.

As shown in **Fig. 2**, we did not identify any significant main effects of cue side (left vs. right) or diagnostic group (TD vs. ASD), nor any Cue side × Group interactions, across any behavioral measures (hit rate, false-alarm rate, d′, or IES). A non-significant Group × Cue-side interaction was observed for RT (F = 1.85, p = 0.174), although the pattern suggested a larger cue-side asymmetry in TD participants, with faster RTs for leftward (582 ms) than rightward cues (596 ms), whereas ASD participants showed minimal differences (615 vs. 617 ms).

**Figure 2.**
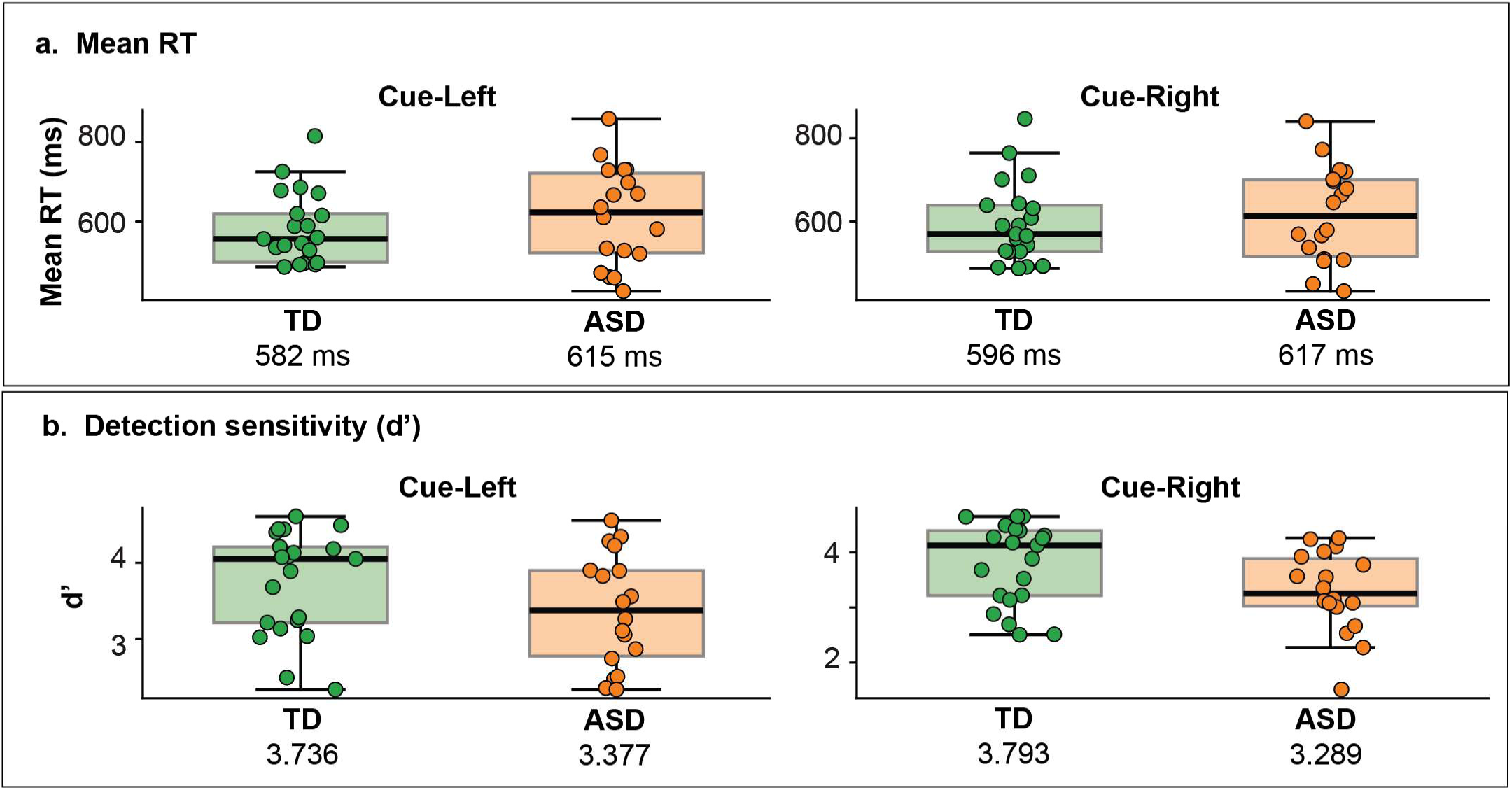
No group differences in behavior. Behavioral performance. Per-participant task performance, separated by group (TD: green; ASD: orange) is shown as **(a)** mean RT (RT) and **(b)** detection sensitivity (d’), plotted separately for Cue Left (left) and Cue Right (right) trials, collapsed across stimulus category and block type. Each dot represents RT for a single participant; boxplots indicate the group distribution (median and interquartile range). The group mean for each measure and cue side is displayed beneath each boxplot.

### S2-Evoked Pupil Response

For peak pupil amplitude to unilateral non-targets, linear mixed-effects analyses revealed a significant main effect of Cue Side (F = 11.45, p = 0.001), with larger pupil responses for stimuli presented during Cue-Right relative to Cue-Left trials (left − right: B = −6.79, p = 0.001; **Fig. 3**; **Supplementary Table 2**). Here, “left” and “right” refer to the cued direction of attention, not the physical location of the stimulus.

**Figure 3.**
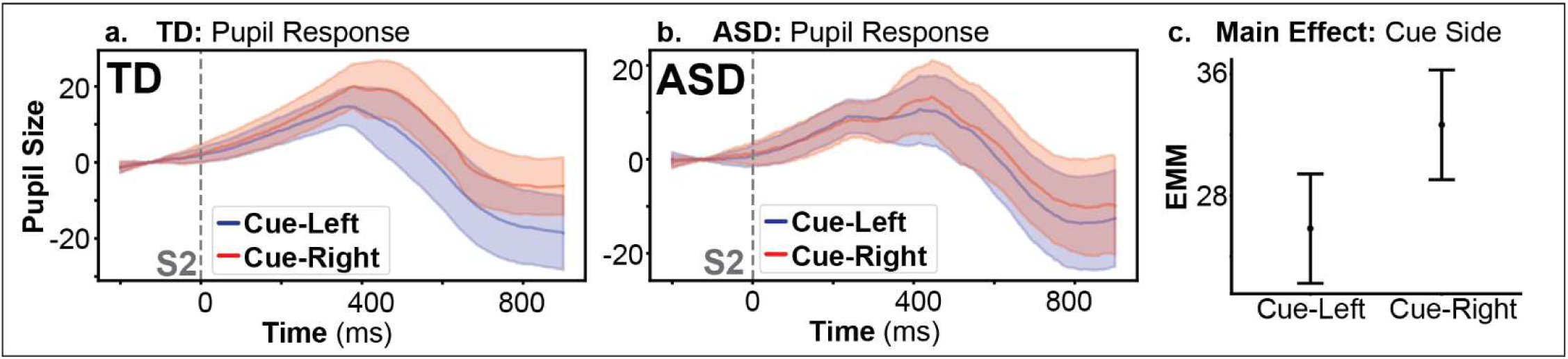
Cue-side effects on the stimulus-evoked pupil response. **a-b.** Grand-average stimulus-evoked pupil responses to unilateral non-target stimuli are shown for Cue Left (blue) and Cue Right (red) trials, separately for TD **(a)** and ASD **(b)** groups, collapsed across pure and intermixed blocks. The solid lines represent the group mean, with shaded regions indicating ±SEM. Time 0 (dashed vertical line) denotes S2 onset. For visualization here, all unilateral stimulus presentations are collapsed regardless of whether a stimulus was presented to the to-be attended or to-be-ignored location, as no significant attentional modulation of pupil response was observed. **c.** Estimated marginal means (EMM) from the linear mixed-effects model revealed a significant main effect of Cue Side. Values are collapsed across group, attentional status and stimulus type. Points represent model-adjusted means, and error bars denote ±1 standard error.

A significant three-way interaction between Cue Side × Attention × Condition was also observed (F = 4.31, p = 0.039). Follow-up analyses indicated significant left–right differences under two specific combinations: when stimuli were unattended in the social condition (left − right: B = −9.07, p = 0.024), and when stimuli were attended in the nonsocial condition (left − right: B = −12.85, p = 0.002), both reflecting larger right relative to left pupil responses. All other post-hoc comparisons were not significant.

No significant main effects or interactions involving group were observed, and there were no significant effects of age.

### S2-Evoked Neural Response

Evaluation of the S2-evoked neural response to non-target unilateral stimuli was conducted using LMMs (see **Supplementary Table 3).**

A robust main effect of Attention was observed on the P1 response. For amplitude, attended stimuli elicited larger responses than unattended stimuli (F = 29.27, p < 0.001; attend − unattend: B = 1.35, p < 0.001). This attention effect was qualified by a significant Cue-Side × Attention × Group interaction for amplitude (F = 4.17, p = 0.042; **Fig. 4c-d**), indicating that attention effects differed by cue direction and diagnostic group. In the ASD group, attended stimuli elicited larger amplitudes than unattended stimuli for leftward (attend − unattend: B = 1.456, p = 0.005), but not rightward (attend − unattend: B = 0.70, p = 0.181) attention. In the TD group, significant attention effects were observed for both leftward (attend − unattend: B = 0.978, p = 0.04) and rightward (attend − unattend: B = 2.259, p < 0.001) attention.

**Figure 4.**
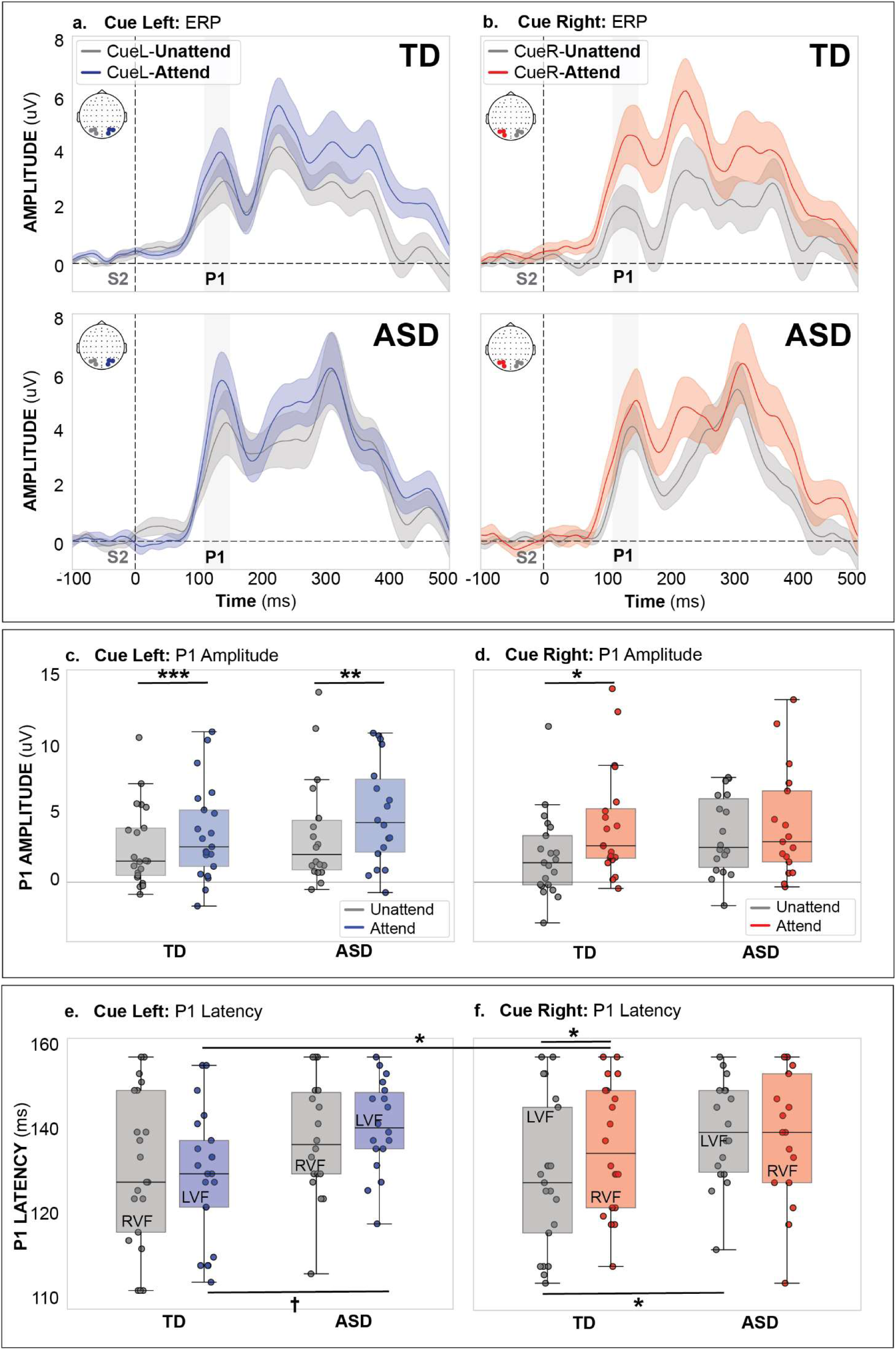
Group differences in timing and amplitude of the stimulus-evoked response. **a-b.** Grand-average event-related potentials (ERPs) elicited by unilateral non-targets are shown for Cue Left **(a)** and Cue Right **(b)** trials, collapsed across pure and inter-mixed blocks, separated by TD (top) and ASD (bottom) groups. Attended (blue: cue-left; red: cue-right) and unattended (gray) waveforms are overlaid, with shaded regions representing ±SEM. Attended responses correspond to stimuli presented in the cued hemifield, whereas unattended responses correspond to stimuli in the uncued hemifield. Time 0 denotes S2 onset (dashed vertical line). The P1 component, indexing early sensory attentional gain, is highlighted by the shaded gray time window (110–150 ms). **c-d.** Boxplots beneath ERP panel summarize mean P1 amplitude elicited by unilateral non-targets, separately for cue-left **(c)** and cue-right **(d)** conditions to compare attended (blue: cue-left; red: cue-right) and unattended (gray) trials. Asterisks indicate significant attention (attended vs. unattended) effects (*p < 0.05, **p < 0.01, ***p < 0.001). **e-f.** Boxplots beneath ERP panel summarize P1 latency elicited by unilateral non-targets, separately for cue-left **(e)** and cue-right **(f)** conditions to compare attended (blue: cue-left; red: cue-right) and unattended (gray) trials. Asterisks indicate significant attention (attended vs. unattended) effects († p<0.1, *p < 0.05, **p < 0.01, ***p < 0.001). LVF: left-visual field; RVF: right visual-field

For latency of the P1 component, a significant Cue-Side × Attention × Group interaction was also observed (F = 4.28, p = 0.04; **Fig. 4e-f**). Within the TD group, attended stimuli during leftward (left visual-field) attention elicited earlier latencies than attended stimuli during rightward (right visual-field) attention (left − right: B = −0.007, p = 0.015). During rightward attention only, unattended (left visual-field) stimuli elicited earlier latencies than attended (right visual-field) stimuli (attend − unattend: B = 0.008, p = 0.01). The ASD group did not exhibit a significant difference in latency for right vs. leftward stimuli, irrespective of attention condition. Additionally, group differences emerged under specific conditions: for leftward attention, ASD participants showed longer latencies to attended (left visual-field) stimuli than TD participants (ASD – TD: B = 0.009, p = 0.035), with a trend in the same direction for rightward (left visual-field) unattended trials (ASD – TD: B = 0.009, p = 0.053). Together, these findings suggest that the lateralized asymmetry observed in TD participants—characterized by relatively faster P1 responses to left visual-field stimuli and consistent with a leftward processing advantage—is not evident in ASD. Instead, ASD participants show reduced lateral differentiation and relatively prolonged responses to left visual-field stimuli.

A significant main effect of Condition was observed for both amplitude and latency. P1 amplitude was reduced in the social relative to nonsocial condition (F = 6.09, p = 0.014; social – nonsocial: B = −0.61, p = 0.014), and P1 latency was earlier in the social relative to nonsocial condition (F = 45.97, p < 0.001; social – nonsocial: B = −0.011, p < 0.001).

Finally, age significantly predicted both P1 measures: increasing age was associated with reduced amplitude (B = −0.564, p < 0.001) and shorter latency (B = −0.002, p = 0.002). No other significant lower-order interactions were observed.

To examine the relationship between early sensory gain and attentional behavior, we assessed the association between P1 amplitude and CPT-3 response style (**Fig. 5**). In the ASD group, reduced P1 amplitude was associated with lower CPT-3 response style scores, indicative of a more impulsive response style. This relationship was observed for both leftward attention (r = 0.56, p = 0.023) and rightward (r = 0.56, p = 0.023) attention. No significant relationship was observed in the TD group (p > 0.05).

**Figure 5.**
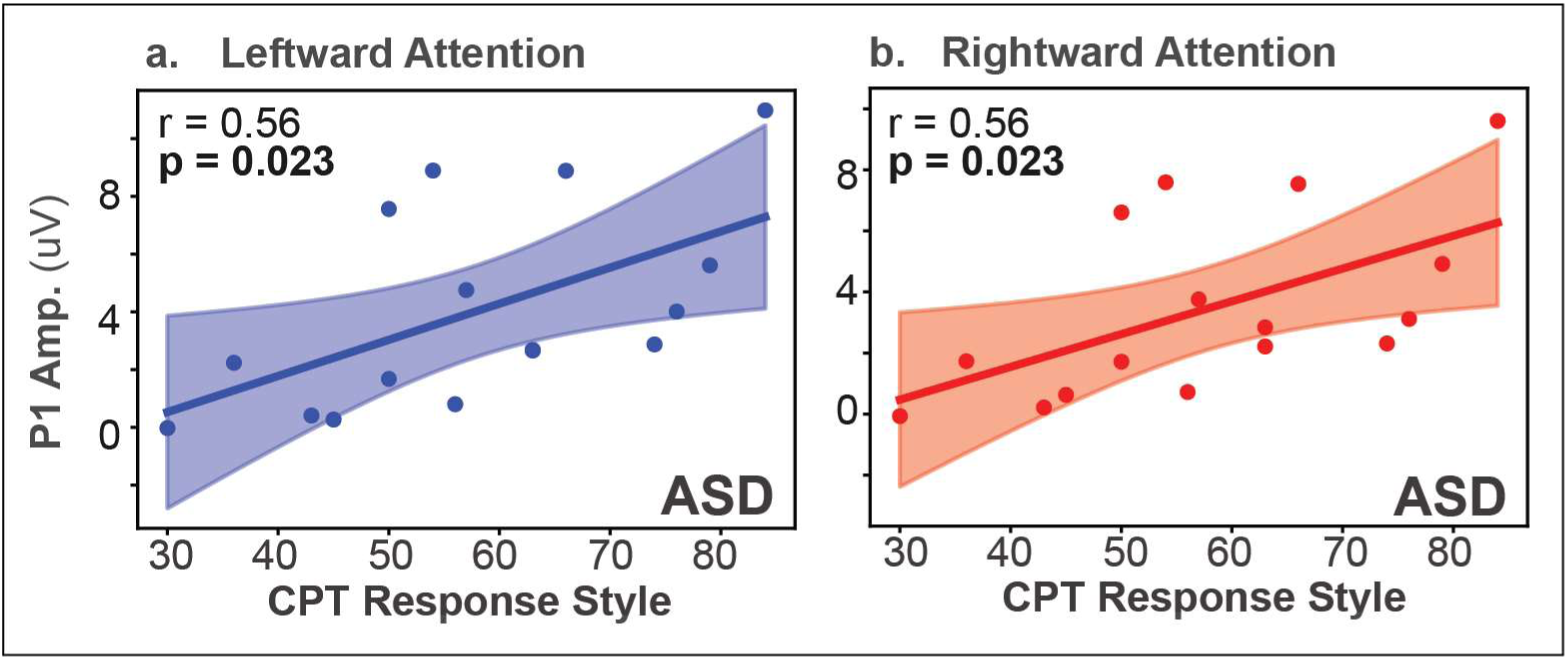
Reduced sensory response in ASD corresponds with a more impulsive clinical attention response style. Relationship between CPT-3 Response Style and P1 amplitude. P1 amplitude was extracted from parieto-occipital electrodes during the 110-150 ms post-stimulus interval, indexing early sensory gain. Each point represents an individual participant; the solid line denotes the best-fitting linear regression, with shaded regions indicating the 95% confidence interval.

### Preparatory Alpha Power

As described in **Supplementary Table 4**, analyses of alpha-power in the anticipatory window revealed a significant main effect of Attention (F = 28.974, p < 0.001), such that alpha power was greater at the unattended relative to attended ROI (unattended − attended: B = 0.273, p < 0.001). This effect was qualified by a significant Attention x Cue-Side interaction (F = 10.058, p = 0.002). Across groups, attentional differences in alpha power were evident for leftward (unattended − attended: B = 0.434, p < 0.001), but not for rightward (B = 0.112, p = 0.119) attention, indicating stronger attentional alpha modulation during leftward attention. Moreover, alpha lateralization differed for the attended versus the unattended ROI: at the attended ROI (RH for cue-left, LH for cue-right), leftward attention elicited lower alpha power (desynchronization) than during rightward attention (left – right: B = −0.251, p = 0.001), whereas no cue-side differences were observed at the unattended ROI.

This pattern was further qualified by a significant Attention × Cue-Side × Group interaction (F = 4.066, p = 0.044 **Fig. 6d-e**), indicating group differences in the spatial organization of alpha modulation. Both groups demonstrated robust attentional modulation of alpha power during leftward attention (unattended – attended; TD: B = 0.535, p < 0.001; ASD: B = 0.332, p = 0.002). During rightward attention, significant attentional modulation was observed only in the ASD group (unattended – attended: B = 0.215, p = 0.043), with no corresponding effect in TD.

**Figure 6.**
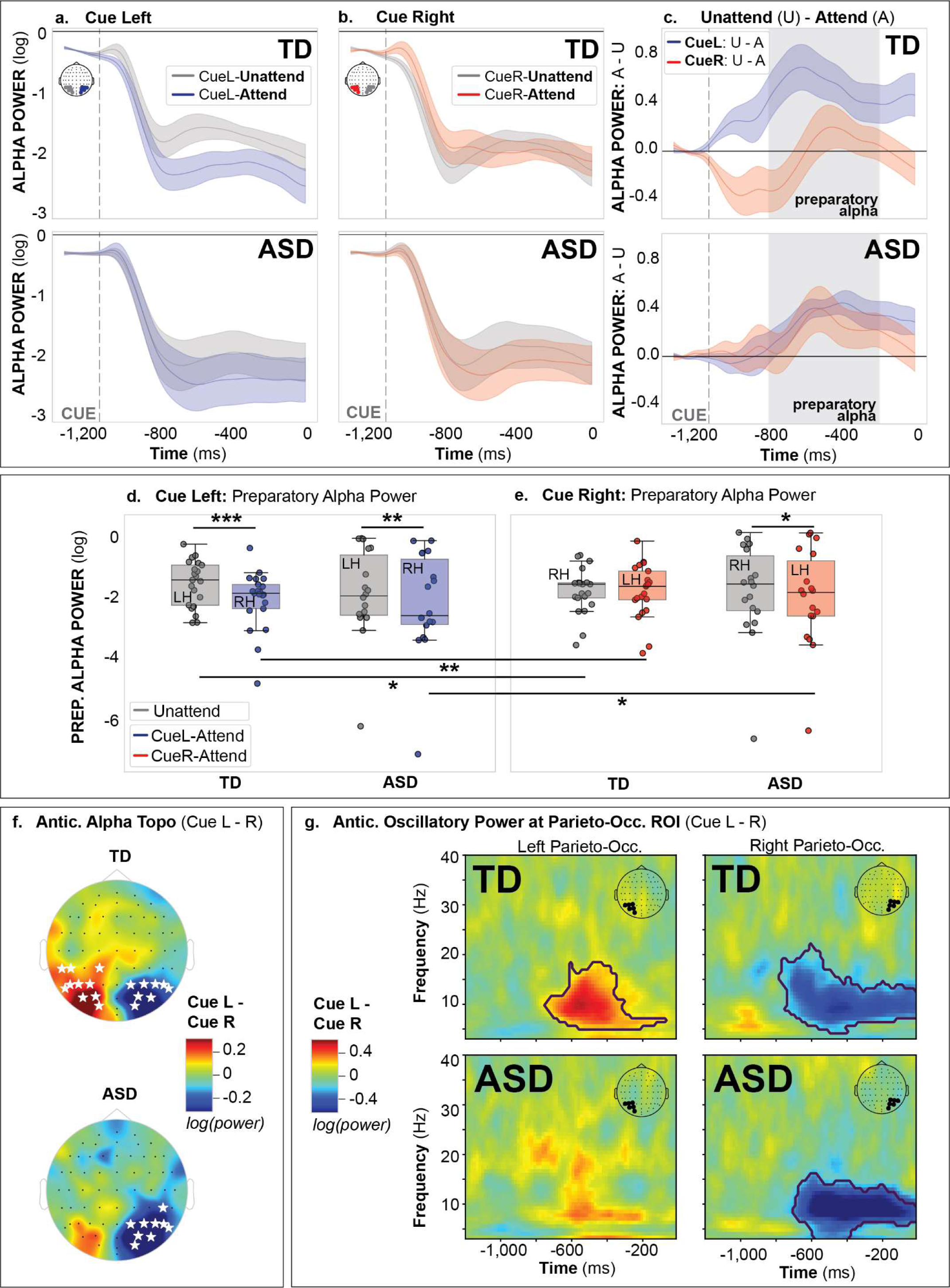
Attentional modulation of alpha-power differs by cue side and group. **a-b.** Time courses of log-transformed alpha-band (7-13 Hz; posterior electrodes) power aligned to cue onset for Cue Left **(a)** and Cue Right **(b)** trials, separated by group (TD: top; ASD: bottom). Attended (blue: cue-left; red: cue-right) and unattended (gray) traces are overlaid, with shaded regions representing ±SEM. Attended responses corresponds to the ROI contralateral to the cued hemifield, and unattended corresponds to the ROI ipsilateral to the cued hemifield.The time course represents the post-cue window prior to S2, in which the dashed vertical line marks cue onset. The preparatory region, indexing preparation for stimuli, is highlighted by the shaded gray time window (−800 to −200 ms). **c.** Attention effects are shown as the difference wave between attended and unattended ROI (A-U) for both Cue Left (blue) and Cue Right (red), separated by group (TD: top; ASD: bottom). **d-e.** Boxplots beneath each ERP panel summarize mean alpha power for Cue Left **(d)** and Cue Right **(e)** trials, separated by group (TD, ASD). Asterisks indicate significant effects of attention (attended vs. unattended ROI) or cue side (left vs. right) (*p < 0.05, **p < 0.01, ***p < 0.001). **f.** Scalp topographies depict preparatory alpha power (7-13-Hz) differences (Cue-Left − Cue-Right), shown separately for TD and ASD groups. Warm colors indicate greater alpha power during cue-left relative to cue-right, observed predominantly over the left hemisphere (which corresponds to the unattended ROI during cue-left and the attended ROI during cue-right). Cool colors indicate the opposite pattern (greater alpha during cue-right). White stars denote electrodes exhibiting significant cue-side differences in alpha power (cluster-based permutation testing, p < .05). **g.** Time–frequency plots (2–40 Hz) depict power differences (Cue-Left − Cue-Right) at parieto-occipital channels (left hemisphere shown on the left; right hemisphere on the right), presented separately for TD (top row) and ASD (bottom row) groups. Warm colors indicate greater power during cue-left relative to cue-right, most prominently over left parieto-occipital regions (corresponding to the unattended ROI during cue-left and the attended ROI during cue-right). Cool colors indicate the opposite pattern (greater power during cue-right). Black contours denote frequencies and time windows exhibiting significant cue-side differences in power (cluster-based permutation testing, p < .05).

The TD group showed significant cue-side modulation of alpha power at both attended and unattended ROIs. At the unattended ROI, leftward attention elicited greater alpha power than rightward attention (left – right: B = 0.244, p = 0.012), whereas at the attended ROI, leftward attention elicited lower alpha power than rightward attention (left – right: B = −0.282, p = 0.004), reflecting a clear bilateral modulation of alpha power (**Fig. 5d-e**). In contrast, the ASD group exhibited cue-side modulation only at the attended ROI, such that leftward attention elicited lower alpha power than rightward attention (left − right: B = −0.22, p = 0.039; **Fig. 6d-e**), whereas in contrast to TD, there was no significant cue-side modulation at the unattended ROI.

Preparatory-alpha lateralization was further characterized by examining cue-dependent hemispheric differences in alpha power (Cue-Left − Cue-Right; **Fig. 6f-g**). The same effects are shown as direct left-versus-right hemispheric comparisons in **Supplementary Fig. 3**. In TD participants, significant cue-side differences were observed bilaterally over parieto-occipital electrodes, with greater alpha power over the left hemisphere during cue-left trials and greater alpha over the right hemisphere during cue-right trials, consistent with spatially specific modulation of posterior alpha during attention (cluster-permutation, *p* < .05). In contrast, ASD participants showed cue-dependent alpha lateralization only over the right parieto-occipital cortex. Time–frequency analysis at parieto-occipital ROIs further confirmed that these effects in TD were driven primarily by sustained modulation within the alpha band (∼7–13 Hz) during the preparatory interval (**Fig. 6g**).

We also identified a significant main effect of Condition (F = 9.235, p < 0.001). Relative to both the Social and Nonsocial conditions, the Intermixed condition was associated with reduced alpha power (Intermixed − Social: B = −0.235, p = 0.001; Intermixed − Nonsocial: B = −0.237, p = 0.001).

No significant effects of Age were observed for anticipatory alpha power.

### Relationship Between Anticipatory Alpha Activity and P1 Amplitude

To assess the relative contributions of anticipatory alpha activity and early sensory responses to behavioral performance, we fit linear mixed-effects models predicting single-trial reaction time (RT) with participant-level random intercepts (**Figure 7**).

**Figure 7.**
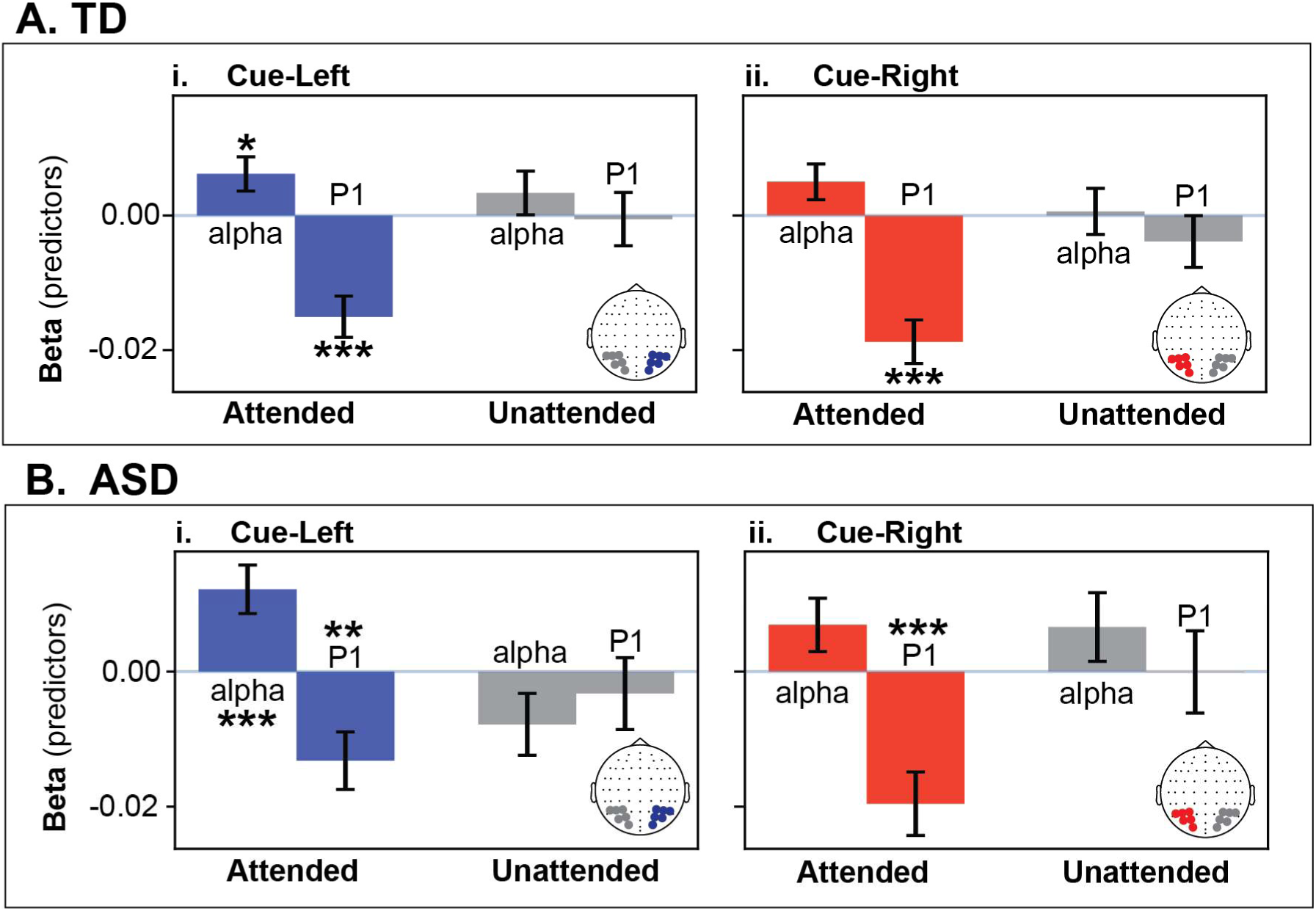
Trial-level oscillatory and sensory predictors of RT. Standardized beta coefficients (± SE) from LMM estimating trial-level RT (RT) as a function of pre-stimulus parieto-occipital alpha power (−800 to −200 ms prior to S2 onset) and P1 amplitude, shown separately for cue-left **(i)** and cue-right **(ii)** trials and for attended (cue-left: right hemisphere; cue-right: left hemisphere) and unattended (cue-left: left hemisphere; cue-right: right hemisphere) ROI. Models are tested separately per group (**A:** TD; **B:** ASD). Positive beta values indicate longer RTs with increasing predictor magnitude, whereas negative beta values indicate faster RTs. Models included subject-level random intercepts to account for within-subject trial clustering. Significance thresholds: * p < .05, ** p < .01, *** p < .001.

Behavioral variability was selectively associated with neural activity over the task-relevant hemisphere. During cue-left trials, reduced pre-target alpha power within the attended ROI was associated with faster responses in both TD (β = 0.0062, SE = 0.0025, p = 0.015) and ASD participants (β = 0.0122, SE = 0.0036, p < 0.001), suggesting that stronger anticipatory alpha power facilitated quicker response times. Concurrently, greater attended P1 amplitudes were linked to shorter RTs in both groups (TD: β = −0.0151, SE = 0.0031, p < .001; ASD: β = −0.0132, SE = 0.0043, p = .002).

A similar relationship was observed for cue-right trials with respect to P1 amplitude. Larger attended P1 responses predicted faster RTs in both TD (β = −0.0187, SE = 0.0032, p < .001) and ASD participants (β = −0.0196, SE = 0.0047, p < .001). However, associations between alpha power and RT did not reach statistical significance in either group (p > 0.05).

To further explore the relationship between anticipatory alpha power and sensory gain of the evoked response in both groups, we next examined whether P1 amplitude covaried with pre-stimulus alpha power at the trial level (**Supplementary Fig. 4**).

In the TD group, we observed a significant, bilateral association between pre-stimulus alpha power and subsequent P1 amplitude over the attended (r = 0.027, p = 0.004) and unattended (r = 0.020, p = 0.029) ROI during leftward attention (**Supplementary Fig. 4A.i**), such that lower alpha states were associated with smaller P1 responses. During rightward attention (**Supplementary Fig. 4A.ii**), where the TD group failed to exhibit significant alpha modulation (**Fig. 5**), neither the attended (LH) nor unattended (RH) ROI demonstrated a significant association between anticipatory alpha power and early sensory gain, as indexed by P1 amplitude (p>0.05).

The ASD group, however, only exhibited a significant relationship between alpha power and P1 amplitude over the right-hemisphere, irrespective of attention or cue side condition. Lower alpha states correlated with smaller P1 responses over the right hemisphere, both during leftward attention when representing the attended ROI (r = 0.032, p = 0.002; **Supplementary Fig. 4B.i**), and during rightward attention when representing the unattended ROI (r = 0.023, p = 0.038; **Supplementary Fig. 4BA.ii**). In the ASD group, the left-hemisphere failed to exhibit a significant association between alpha power and P1 amplitude in any condition (p>0.05).

### Resting State

In an analysis of periodic alpha-band amplitude at rest, we failed to identify main effects of Group or Hemisphere. The main effect of Condition (F = 858.473, p < 0.001) indicated greater alpha amplitude during the eyes-closed relative to eyes-open state (open − closed: B = −0.356, p < 0.001). This was qualified by a Condition × Group interaction (F = 4.300, p = 0.038), reflecting a numerically stronger, though not significantly different (p > 0.05), eyes-closed enhancement in ASD (open − closed: B = −0.381, p < 0.001) compared with TD (open − closed: B = −0.331, p < 0.001). A significant Condition × RS-block interaction (F = 114.078, p < 0.001) indicated that alpha amplitude was greater during eyes-closed than eyes-open both before (open − closed: B = −0.486, p < 0.001) and after (open − closed: B = −0.226, p < 0.001) the task. However, task-related changes differed by condition: eyes-open alpha amplitude was lower before compared to after the task (before – after: B = −0.107, p < 0.001), whereas eyes-closed alpha amplitude decreased from before to after the task (before – after: B = 0.152, p < 0.001). This pattern was further modulated by group (Condition × Group × RS-block), such that the pre- to post-task increase in eyes-open alpha amplitude was observed in ASD (before – after: B = −0.197, p < 0.001) but not in TD (before – after: B = −0.035, p = 0.135). In contrast, both groups exhibited a reduction in eyes-closed alpha amplitude from before to after the task (ASD before – after: B = 0.151, p < 0.001; TD before – after: B = 0.154, p < 0.001). Finally, a RS-block × Group interaction (F = 9.099, p = 0.003) indicated greater alpha amplitude before versus after the task in TD only (before – after: B = 0.059, p < 0.001), with no main effects of RS-block or age.

We also evaluated aperiodic activity at rest to probe E/I integrity. No main effects of Group (**Fig. 8a**) or Hemisphere were observed; however, a significant Hemisphere × Group interaction (F = 20.253, p < 0.001) indicated a marginally greater aperiodic exponent in the right relative to left parieto-occipital ROI in ASD only (left – right: B = −0.061, p = 0.079; **Fig. 8b**). The main effect of Condition (F = 193.821, p < 0.001) revealed higher aperiodic exponent during the eyes-closed relative to eyes-open state (open − closed: B = −0.136, p < 0.001). A main effect of RS-Block (before vs. after task) was also observed (F = 135.939, p < 0.001), reflecting higher exponent prior to compared with following the task (before − after: B = 0.114, p < 0.001). These effects were qualified by a significant Condition × RS-Block interaction (F = 4.189, p = 0.041), such that aperiodic exponent was higher in the eyes-closed relative to eyes-open condition both before (open − closed: B = −0.116, p < 0.001) and after (open − closed: B = −0.156, p < 0.001) the task. Additionally, both eyes-open (before − after: B = 0.134, p < 0.001) and eyes-closed (before − after: B = 0.094, p < 0.001) conditions showed higher exponent before relative to after the task. A RS-Block × Group interaction (F = 5.255, p = 0.022) further indicated greater pre-versus post-task exponent in both TD (before − after: B = 0.136, p < 0.001) and ASD (before − after: B = 0.091, p < 0.001), with no significant difference between groups in post hoc comparisons (p > 0.05). Finally, a main effect of Age was observed (B = −0.042, p = 0.002), indicating lower aperiodic exponent in older participants.

**Figure 8.**
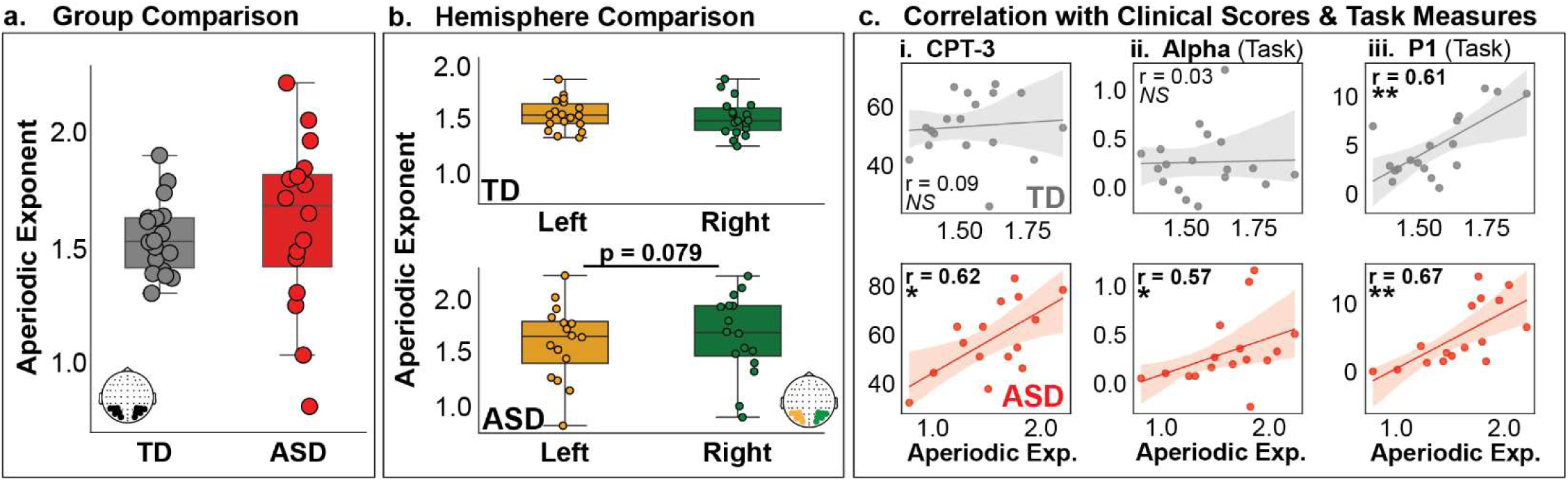
Group differences, hemispheric effects, and clinical correlations of aperiodic exponent at rest. **a.** Boxplots show the distribution of aperiodic exponent values over parieto-occipital region (collapsed over left and right ROI) in typically developing (TD; grey) and autism spectrum disorder (ASD; red) participants. Individual data points are overlaid. **b.** Aperiodic exponent is plotted separately for left and right parieto-occipital ROIs within each group, shown for TD (top) and ASD (bottom) participants. The reported p-value reflects the difference in resting-state aperiodic exponent between the left and right hemispheres. **c.** Scatterplots illustrate the relationship between aperiodic exponent and **(i)** CPT-3 response style, **(ii)** preparatory alpha-band power, and **(iii)** P1 peak amplitude, shown separately for TD (grey; top) and ASD (red; bottom) participants. CPT-3 scores index response style during a button-press task as a clinical measure of attention, with lower scores reflecting a more impulsive response style, 50 indicating normative performance, and higher scores reflecting a more cautious response style. Task-related alpha power was quantified using Morlet wavelet decomposition in the −800 to −200 ms pre-stimulus interval (S2 preparatory period) and averaged across attended and unattended parieto-occipital ROIs and task conditions, providing an index of the strength of anticipatory oscillatory activity. P1 peak amplitude was defined as the maximum positive deflection within the canonical P1 time window following stimulus onset for unilateral non-target trials, indexing early sensory gain, and was averaged across ROIs and conditions. Regression lines with shaded 95% confidence intervals are shown. Reported p-values reflect the strength of association based on Pearson or Spearman correlation analyses, as determined by data normality. NS = not significant; *\*p* < 0.05; \*\**p* < 0.01; \*\*\**p* < 0.001.

We next examined relationships between aperiodic exponent and clinical attention, task-related alpha power, and early sensory responses (**Fig. 8c**). In TD participants, no significant associations were observed between aperiodic exponent and CPT-3 scores (r = 0.09, p = 0.729) or task-related alpha power (r = 0.03, p = 0.911). However, the aperiodic exponent was positively associated with P1 amplitude (r = 0.61, p = 0.007), indicating that steeper aperiodic slopes were linked to enhanced early sensory gain. In contrast, ASD participants exhibited significant positive associations across all measures. The aperiodic exponent positively correlated with CPT-3 scores (r = 0.62, p = 0.0140), reflecting a relationship between flatter aperiodic slope and a more impulsive response style. Similarly, the aperiodic exponent at rest was positively associated with task-related anticipatory alpha power (r = 0.57, p = 0.022) and P1 amplitude (r = 0.67, p = 0.004). Aperiodic exponent was not associated with behavioral performance (RT, d’, IES, hit rate, FA rate) for TD or ASD groups (p>0.05).

## Discussion

### Similar Cortical Mechanisms of Leftward Spatial Attention in ASD and TD

Leftward attention appeared largely intact in ASD across all measures. Behaviorally, TD and ASD participants showed comparable performance when responding to targets in the left visual field. At the neural level, both groups exhibited marked similarity in P1 amplitude modulation and alpha-band dynamics, indicating comparable sensory gain mechanisms supporting spatial attention during leftward attention. Task-dependent modulation of both alpha power (Worden, Foxe et al. 2000, Kelly, Gomez-Ramirez et al. 2009) and P1 amplitude (Luck, Heinze et al. 1990, Heinze, Mangun et al. 1994, Wijers, Lange et al. 1997, Woldorff, Fox et al. 1997) is a well-established feature of visuospatial attention in typically developing populations; however, these neural mechanisms remain fundamentally understudied in ASD.

Furthermore, in both groups, alpha modulation was coupled to P1 amplitude over the attended ROI, suggesting coordinated interactions between preparatory oscillatory dynamics and early sensory amplification during leftward attention. In addition, reduced alpha power and enhanced P1 amplitude over the attended ROI predicted faster reaction times in both groups, demonstrating the behavioral relevance of these neural mechanisms for speeded target detection. Previous work offers supporting evidence of a robust relationship between anticipatory-alpha power and task-related performance (Thut, Nietzel et al. 2006, Kelly, Gomez-Ramirez et al. 2009, Murphy, Foxe et al. 2014), though this relationship is poorly understood in autism. Moreover, the relationship between anticipatory alpha activity and P1 amplitude remains an ongoing debate in the literature, with some proposing that the P1 primarily reflects ongoing alpha phase dynamics (Gruber, Klimesch et al. 2005, Klimesch, Hanslmayr et al. 2007, Fellinger, Klimesch et al. 2011, Klimesch, Fellinger et al. 2011, Trajkovic, Di Gregorio et al. 2024), while others argue that these processes are functionally dissociable (Slagter, Prinssen et al. 2016). The coupling observed here, particularly given its relationship to behavioral performance, provides new insight into how preparatory oscillatory states and early sensory gain mechanisms interact during spatial attention.

Overall, these findings suggest that the core attentional control functional architecture supporting leftward visuospatial selective attention is largely similar in ASD and TD, with robust coordination between posterior oscillatory dynamics, early sensory gain mechanisms and behavior.

### Differing Cortical Mechanisms of Rightward Spatial Attention in ASD and TD

In contrast, clear group differences in the neural mechanisms underlying rightward attention were observed. In the TD group, cue-right trials elicited enhanced P1 sensory responses to attended stimuli, whereas ASD participants failed to show comparable amplification of early visual processing. Furthermore, preparatory oscillatory dynamics also diverged between groups. As we showed previously (Darrell, Vanneau et al. 2026), TD participants only exhibited preparatory alpha modulation during leftward and not during rightward attention, showing hemispheric differences in how spatial attention is deployed. In contrast, ASD participants showed preparatory alpha modulation during rightward—in addition to during leftward—attention, suggesting a more uniform oscillatory pattern in ASD, irrespective of cue direction. However, alpha power over the attended region during rightward attention in the ASD cohort was not associated with changes in P1 amplitude, contrasting with the clear alpha–P1 coupling observed for both groups during leftward attention. Moreover, faster reaction times in both groups were selectively associated with increased P1 amplitude over the attended hemisphere, whereas alpha power did not predict behavioral performance. Together, these findings suggest that anticipatory alpha recruitment in ASD during rightward attention may not effectively translate into behaviorally relevant sensory facilitation.

Previous studies have also reported differences in attentional sensory gain in ASD, including increased visual P1 attentional modulation consistent with hyper-focused allocation (in autistic adults with parietal cortex abnormalities) and reduced auditory N1 attentional modulation (Townsend and Courchesne 1994, Teder-Salejarvi, Pierce et al. 2005). However, these findings are based on small adult samples and do not assess spatial asymmetries. Similarly, although some prior work has identified attenuations in stimulus-evoked alpha desynchronization in ASD (Keehn, Nair et al. 2016, Keehn, Westerfield et al. 2017, Canigueral, Palmer et al. 2022), no studies to date have examined preparatory alpha-band mechanisms in autism within a strictly visuospatial attention paradigm. Notably, one previous study reported that ASD participants failed to exhibit posterior alpha-band modulation in an intersensory attention task, suggesting differences in the deployment of alpha-band mechanisms, albeit in a different context (Murphy, Foxe et al. 2014).

Together, the current findings provide the first characterization of cue-specific alterations in the cortical mechanisms underlying spatial attention in autism, revealing dissociable disruptions in anticipatory alpha modulation and P1 sensory gain. Although ASD participants recruited anticipatory alpha dynamics during rightward attention, these oscillatory mechanisms did not appear to effectively translate into downstream sensory facilitation, suggesting disrupted coupling between top-down oscillatory control and early visual processing during rightward orienting. Importantly, the neural differences identified in the present study did not correspond to overt behavioral impairments in ASD, suggesting that the neural mechanisms most directly linked to behavior (namely attended P1 amplitude) remain preserved in this context. Instead, the absence of significant attentional modulation of P1 amplitude in ASD may reflect comparatively elevated responses to unattended stimuli, suggesting reduced suppression of task-irrelevant distractors rather than deficient sensory gain to attended targets per se. Although interpretive, this possibility aligns with broader literature describing hypo-focus and enhanced visual distractor processing in ASD (Remington, Swettenham et al. 2009, Ohta, Yamada et al. 2012, Remington, Swettenham et al. 2012).

### Baseline Cortical Excitability in ASD

To determine whether the task-related neural differences observed here may reflect baseline cortical alterations in autism, we additionally examined resting-state cortical dynamics. These analyses failed to demonstrate significant group-level differences in cortical excitability—as indexed by the aperiodic exponent— or alpha power at rest, suggesting that the neural differences observed during the task are not broadly attributable to baseline differences. However, within-group hemispheric analyses revealed a moderate leftward asymmetry in the ASD group, characterized by a reduced aperiodic exponent in the left hemisphere relative to the right, which was not observed in TD participants. The aperiodic exponent, reflecting the 1/f slope of the power spectrum, indexes the distribution of spectral energy across frequencies (Buzsaki and Draguhn 2004). A reduced exponent corresponds to a flatter spectrum and is often interpreted as relatively increased cortical excitability (corresponding to a shift in excitation–inhibition (E/I) balance), whereas a steeper exponent is associated with reduced excitability (Gao, Peterson et al. 2017, Lombardi, Herrmann et al. 2017, Brake, Duc et al. 2024). Prior work from both animal (Yizhar, Fenno et al. 2011, Juarez and Martinez Cerdeno 2022, Bryers, Hawkes et al. 2024) and human (Bozzi, Provenzano et al. 2018, Port, Oberman et al. 2019, Bertelsen N 2023, Huang, Chen et al. 2026) work has reported alterations in excitation–inhibition balance in ASD (Cellot and Cherubini 2014, Uzunova, Pallanti et al. 2016), although findings remain mixed (Edgar, Fisk et al. 2016, Ono, Kudoh et al. 2020, Ahlfors, Graham et al. 2024, Darrell, Vanneau et al. 2025), suggesting that these observed differences are heterogeneous and may be either i) localized to specific circuits and/or ii) present at specific developmental stages, rather than reflecting a global and developmentally stable imbalance.

In the present study, across both groups, a lower resting-state aperiodic exponent was associated with reduced P1 amplitude during task performance. In the ASD group specifically, a lower resting-state aperiodic exponent (and lower P1 amplitude) was also linked to lower CPT-3 response style scores, independent of attentional direction. The CPT-3 response style metric reflects the speed–accuracy tradeoff during a standardized and normed clinical attention task, with lower scores indicating a more impulsive response bias that prioritizes speed over accuracy. Together, these findings indicate that baseline alterations in cortical dynamics at rest may have implications for task-evoked sensory processing and may relate to clinically relevant attentional behavior in ASD. This pattern aligns with prior work linking altered aperiodic activity at rest (Shuffrey, Pini et al. 2022, Arutiunian, Santhosh et al. 2025, Chung, Job Said et al. 2025, Wilkinson, Chung et al. 2025) to restricted and repetitive behaviors (Chung, Job Said et al. 2025) and language delay (Wilkinson, Chung et al. 2025) in infants at risk for autism and in autistic children.

It is possible that altered left-hemisphere (LH) resting-state dynamics may also contribute to the task-related differences observed over the LH in ASD, including the absence of typical task-dependent modulation of alpha activity (**Fig. 6f–g**) and the decoupling of alpha–P1 interactions across both attentional conditions (**Supplementary Fig. 4**) observed over the LH. These observations are broadly supported by literature spanning diverse clinical cohorts and cognitive and sensory domains that report altered LH function in ASD (Eyler, Pierce et al. 2012, Fiebelkorn, Foxe et al. 2013, Perkins, Stokes et al. 2014, Peterson, Mahajan et al. 2015, Stroganova, Komarov et al. 2020), including during visuospatial attention tasks (Belmonte and Yurgelun-Todd 2003). In this context, the present findings suggest that LH mechanisms may play a particularly important role during rightward attention. This interpretation may be further understood within the broader framework of right-hemisphere dominance for visuospatial attention, which proposes that the right hemisphere contributes to attentional orienting across both hemifields, whereas the left hemisphere is more selectively involved in rightward orienting (Mesulam 1999, Corbetta, Kincade et al. 2000, Foxe, McCourt et al. 2003). Consistent with this framework, we show faster reaction times during rightward attention were primarily associated with greater target-side (LH) P1 sensory gain, which may help explain why rightward attention appears atypical in ASD, where LH mechanisms may be engaged less effectively. Nevertheless, the present findings remain preliminary, and further investigation is needed to clarify the specific contribution of LH dynamics to attentional processing differences in ASD.

### Limitations and Future Work

A limitation of this study is its relatively small sample size, due in part to use of strict quality control procedures to ensure central fixation compliance, task engagement, and high-quality physiological data. While necessary, the final sample may also introduce sampling bias. Furthermore, the current sample only includes ASD individuals with relatively low support needs. While necessary to accommodate a cognitively demanding experimental paradigm and long data collection session, future studies using shorter or more flexible paradigms (e.g., fixation gated stimulus presentation) are needed to test the generalizability of these findings to a broader clinical population.

Another limitation is that the study design may have limited our ability to detect the influence of social stimuli on spatial attention, an original goal of our study. Although we expected social versus non-social attention to differ in ASD given the hallmark social processing differences associated with the disorder, attentional effects were not influenced by whether face or house stimuli were used. One possible explanation is that the task required participants detect a target ring superimposed on the images, rather than attend to features of the faces or houses themselves. As a result, the paradigm may not have sufficiently engaged social perceptual processing mechanisms. Future studies should incorporate tasks that require explicit processing of facial or object features to more directly test potential differences in spatial attention in the context of social environments in ASD.

## Conclusions

This work offers a systems-level account of how posterior oscillatory dynamics and early sensory gain interact to support spatial attention in autism, addressing a key gap in understanding atypical attention in ASD. Critically, leftward attentional cortical mechanisms appeared strikingly similar in ASD and controls, with preserved behavioral performance, alpha modulation and sensory gain, indicating that core attentional architecture is maintained. Group differences emerged primarily during rightward spatial attention, where canonical sensory gain mechanisms that were present in the control group were diminished in ASD, alongside recruitment of anticipatory alpha-band modulation that was not observed in TD participants. Together, these findings argue against the notion of global spatial attention differences in ASD and instead point to a direction-specific reorganization of attentional control, as demonstrated visually in **Fig. 9**. Importantly, while behavior was preserved in this highly constrained task, everyday environments place substantially greater demands on flexible, dynamic attentional control, suggesting that this altered functional architecture may contribute to the attentional differences commonly observed in autism.

**Figure 9.**
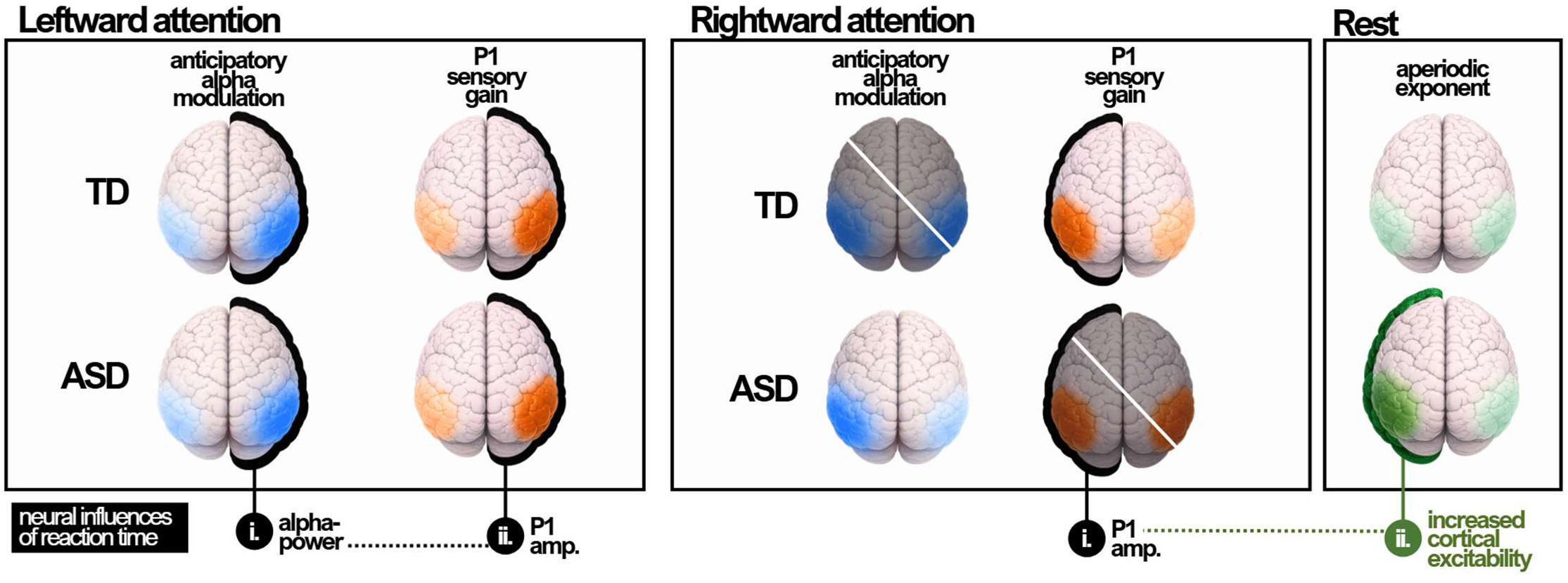
Proposed model of asymmetric attentional control mechanisms in ASD. During leftward attention, both typically developing (TD) and ASD groups exhibited canonical attentional modulation, characterized by anticipatory parieto-occipital alpha modulation and enhanced P1 sensory gain over the attended hemifield. In both groups, anticipatory alpha activity was coupled to P1 sensory gain, and both neural processes significantly influenced reaction time, suggesting relatively preserved leftward attentional control mechanisms in ASD. Rightward attention, however, revealed a dissociation between groups. TD participants showed P1 sensory gain in the absence of significant alpha modulation, whereas ASD participants showed the opposite pattern, anticipatory alpha modulation but no corresponding P1 enhancement. This suggests that although ASD participants recruited alpha-based control mechanisms during rightward attention, this did not translate into effective sensory facilitation. Consistent with this, reaction time was associated with P1 amplitude but not alpha power in both groups, highlighting the behavioral relevance of sensory gain over oscillatory control. Resting-state analyses further revealed moderately asymmetric baseline cortical dynamics in ASD, characterized by a reduced aperiodic exponent over left parieto-occipital cortex, potentially reflecting increased cortical excitability. Lower aperiodic exponent at rest was associated with reduced rightward P1 sensory gain across both groups, suggesting that altered baseline excitability may interfere with the efficient deployment of rightward visuospatial attention, where behavioral performance appears to depend primarily on early sensory gain mechanisms indexed by the P1.

## Author Contributions

M.D and S.M conceived the study. M.D was responsible for participant recruitment and data collection and curation. M.D pre-processed and analyzed the data under the supervision of T.V, C.B and S.M. M.D wrote the first draft of the manuscript. S.M, J.J.F, T.V and C.B provided edits on the manuscript. All authors approved the final manuscript.

## List of Abbreviations

ADOS-2: Autism Diagnostic Observation Schedule, 2nd Edition
ASD: Autism spectrum disorder
CPT-3: Conners’ Continuous Performance Test, 3rd Edition
d′: Detection sensitivity (d-prime)
EEG: Electroencephalography
ERP: Event-related potential
FA: False alarm
FFT: Fast Fourier Transform
FS-IQ: Full-scale intelligence quotient
Hz: Hertz
ICA: Independent component analysis
IES: Inverse efficiency score
IM: Inter-mixed (block condition)
LMM: Linear mixed-effects model
LSL: Lab Streaming Layer
NS: Non-social
PO: Parieto-occipital
P1: First positive visual ERP component (∼100 ms)
ROI: Region of interest
RT: Reaction time
RS: Resting state
S: Social
SEM: Standard error of the mean
SOA: Stimulus-onset asynchrony
SRS-2: Social Responsiveness Scale, 2nd Edition
S1: Stimulus 1 (cue)
S2: Stimulus 2 (stimuli/target display)
TD: Typically developing
UDTR: Up-Down Transformed Rule
V-IQ: Verbal intelligence quotient

## Acknowledgements & Funding

Support for recruitment and phenotyping of participants was provided by the Human Clinical Phenotyping Core of the NICHD funded Rose F. Kennedy Intellectual and Developmental Disabilities Research Center (P50 HD105352). Work at the collaborating site in Rochester is supported through the B. Thomas Golisano Intellectual and Developmental Disabilities Research Institute (UR-IDDRC), which is supported by a center grant from the Eunice Kennedy Shriver National Institute of Child Health and Human Development (P50 HD103536 – to JJF). Partial support was also provided by the Albert Einstein College of Medicine Medical Scientist Training Program (T32-GM149364).

