## Supplementary Materials for "Asymmetric neural dynamics of visuospatial attention in autism spectrum disorder"

### Supplementary Methods

##### *Equating Task Difficulty Across Participants via UDTR*

Prior to the experimental task, participants completed an adaptive staircase calibration designed to equate performance across individuals. This procedure, known as the Up-Down Transformed Rule (UDTR) was used to rapidly equate performance across the task and across participants (Wetherill & Levitt, (Wetherill and Levitt 1965)) before the beginning of the formal experimental sessions. Participants were instructed to respond to the presence of a white target ring superimposed on a face or a house via keyboard press (left key = present; right key = absent). Correct responses triggered an adaptive staircase procedure that adjusted stimulus contrast based on accuracy. We used a 3-up, 1-down rule, meaning that, when a participant made three consecutive correct responses, the stimulus contrast level was decreased by one step, and for any incorrect response, stimulus contrast level was increased by one step. This set of rules converges on an accuracy level of 79.4%.

For this calibration phase, only right-hemifield, unisensory S2 stimuli were presented, following an approach similar to the unisensory thresholding procedure described by Murphy et al. (Murphy, Foxe et al. 2016). Consequently, while providing a common performance starting point for all participants, the resulting contrast thresholds do not incorporate the additional competitive demands introduced by bilateral stimulus presentations in the main task, nor do they capture potential differences in sensitivity between hemifields.

##### *Pre-Processing*

Eye-Tracking

Eye-tracking—which was exported using Data Viewer software Version 4.3.210—served two distinct purposes in the present study: (1) to confirm compliance with covert spatial attention instructions by ensuring central fixation was maintained, and (2) to measure task-evoked pupil responses as an index of arousal and cognitive effort.

*Quality Control*

Eye tracking data were cleaned individually for each participant using custom Python scripts and following best practices (Mathot 2018, Mathot and Vilotijevic 2023). Offscreen data were removed, blinks and artifacts were identified and corrected. After cleaning, trials missing over 50% of pupil data were excluded.

*Identification and Exclusion of Saccade-Contaminated Trials*

Additional gaze-based preprocessing was performed to ensure stable central fixation at stimulus onset. Because gradual drift in eye-tracker calibration can introduce block-specific offsets, the fixation center was estimated separately for each participant and block. For this purpose, gaze samples from a pre-cue baseline window (−1.22 to −1.12 s) were pooled across trials, and the median horizontal and vertical gaze positions were computed to define the block-specific fixation center. This approach allowed robust correction for slow calibration drift or posture-related offsets.

Fixation stability was then evaluated at stimulus onset (−100 to +100 ms). For each trial, gaze samples were classified as within fixation if their two-dimensional distance from the estimated fixation center fell within a circular region of interest with a radius of 2° of visual angle. Trials were retained when at least 80% of valid (non-missing) gaze samples within this window fell inside the fixation region; otherwise, they were excluded.

Participants were excluded if fewer than 50% of trials met this fixation criterion, indicating excessive saccadic behavior incompatible with the covert attention requirements of the task.

EEG

EEG data were analyzed using Python (MNE; (Gramfort, Luessi et al. 2013)) and custom in-house scripts. Raw data were low-pass FIR-filtered at 40 Hz and high-pass FIR-filtered at 0.01 Hz.

Bad channels were detected automatically using *pyPrep.NoisyChannels ((Bigdely-Shamlo, Mullen et al. 2015))*, which includes a combination of classical bad-channel detecting functions: low signal-to-noise ratio, low RANSAC (Martin A. Fischler 1981) prediction or correlation with other channels, abnormal amplitude or amounts of high-frequency noise, and near-flat channels. Subjects were excluded if >15% of channels were identified as bad; otherwise, bad channels were interpolated using the spherical-spline method (Perrin, Pernier et al. 1989).

Epochs were constructed from a -2,280 to +1,000 ms time window surrounding the onset of S2 (t=0 ms). Independent Component Analysis (ICA) using the *fastica* algorithm (max 1,000 iterations) was applied to epoched data high-pass filtered at 1 Hz to identify and manually remove components corresponding to eye movements, including horizontal saccades and blinks. The 1 Hz high-pass filter was applied only for ICA component identification, and all subsequent analyses were conducted on the original data (0.01-40 Hz) after applying ICA removal.

Epochs containing artifacts were excluded by automatic artifact rejection *(autoreject; (Jas, Engemann et al. 2017))*, which utilizes a cross-validation metric to identify an optimal amplitude threshold (Jas, Engemann et al. 2017). Subjects were excluded if >50% of trials were rejected across all conditions.

Epochs were averaged at the individual subject level, for each of the stimulus conditions of interest, for subsequent analyses. Data was re-referenced to the average of all electrodes, a commonly used reference technique for evaluating event-related potentials (Tsuchimoto, Shibusawa et al. 2021).

#### Supplementary Figures

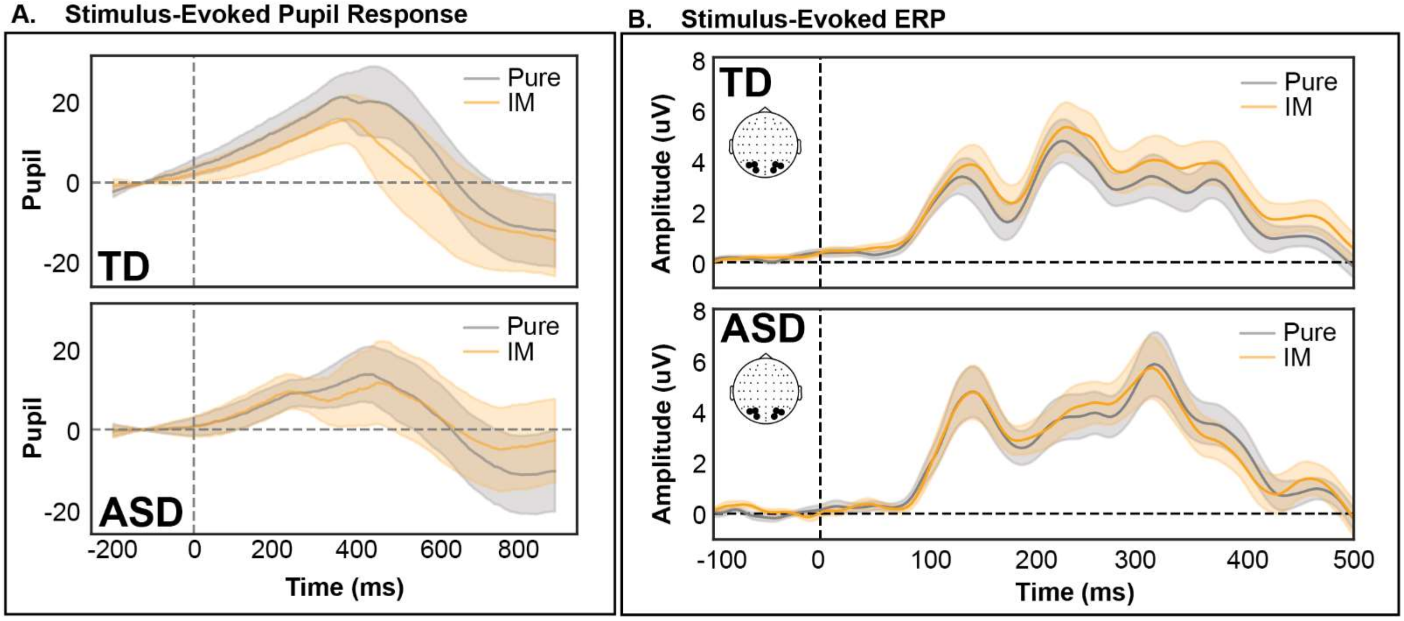


**Supplementary Figure 1. Stimulus-evoked pupil and ERP responses by block type.**
**A.** Grand-average stimulus-evoked pupil responses for pure (grey) and intermixed (IM; orange) blocks are shown separately for TD (top) and ASD (bottom) groups. Shaded regions represent ± SEM across participants. Time 0 ms denotes S2 onset. **B.** Grand-average stimulus-evoked ERPs at contralateral parieto-occipital electrodes are shown for pure (grey) and intermixed (IM; orange) blocks, separately for TD (top) and ASD (bottom) groups. Shaded regions represent ± SEM. To assess potential differences between block types, cluster-based permutation testing was applied to the pure–intermixed difference wave (pure − IM) separately for each group. No significant clusters were observed for either pupil or ERP responses in TD or ASD groups (all $p > 0.05$), supporting the collapse across block types in subsequent analyses.

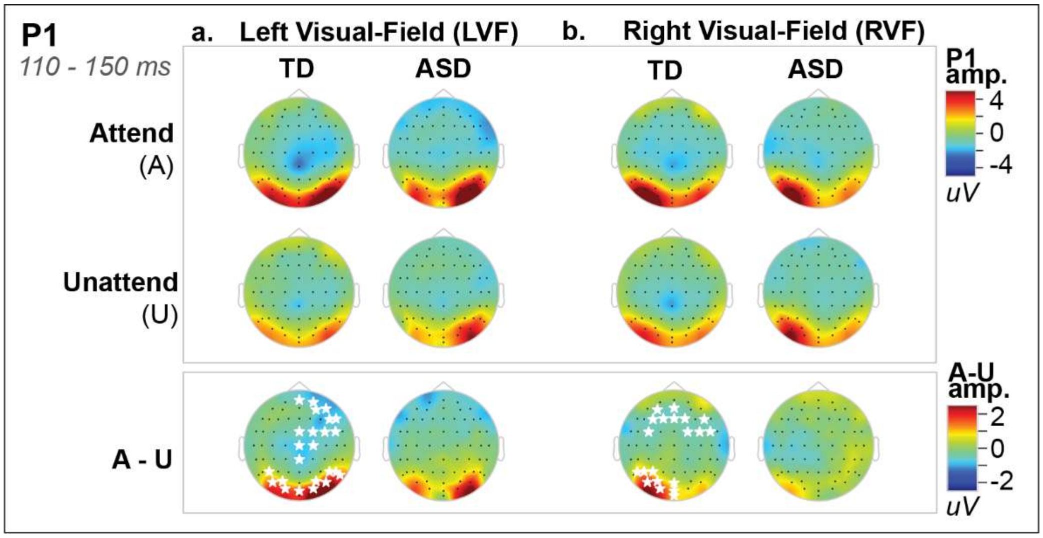


**Supplementary Figure 2. Early P1 amplitude varies as a function of visual field and attention in TD and ASD.**

**a-b.** Topography of P1 amplitude elicited by unilateral non-targets are shown for stimuli presented in the left **(a)** and right **(b)** visual-fields, separated for TD and ASD groups. For each visual field, maps are shown separately for attended (A) and unattended (U) conditions.

For each visual field, attended stimuli refer to trials in which attention was directed toward that hemifield (cue left for LVF; cue right for RVF). Unattended stimuli refer to trials in which attention was directed to the opposite hemifield (cue right for LVF; cue left for RVF).

The bottom row displays the attentional modulation (A − U) of P1 amplitude for each visual field and group. White stars denote electrodes showing significant attentional effects (cluster-permutation testing, $p < .05$).

To characterize the spatial distribution of early sensory gain and determine whether attentional modulation differed as a function of stimulus hemifield, we examined P1 amplitude separately for stimuli presented in the left and right visual fields and computed the attentional difference (A − U) for each group (**Supplementary Fig. 1**). In TD participants, significant attentional enhancement of P1 amplitude was observed for left visual field stimuli over bilateral parieto-occipital electrodes, whereas for right visual field stimuli the effect was restricted to a cluster of electrodes in the left-hemisphere. In ASD participants, P1 attentional modulation for stimuli presented in either visual field did not reach statistical significance using cluster-permutation testing, although the topographies visually suggest bilateral attentional modulation for left visual field stimuli and markedly reduced modulation for right visual field stimuli.

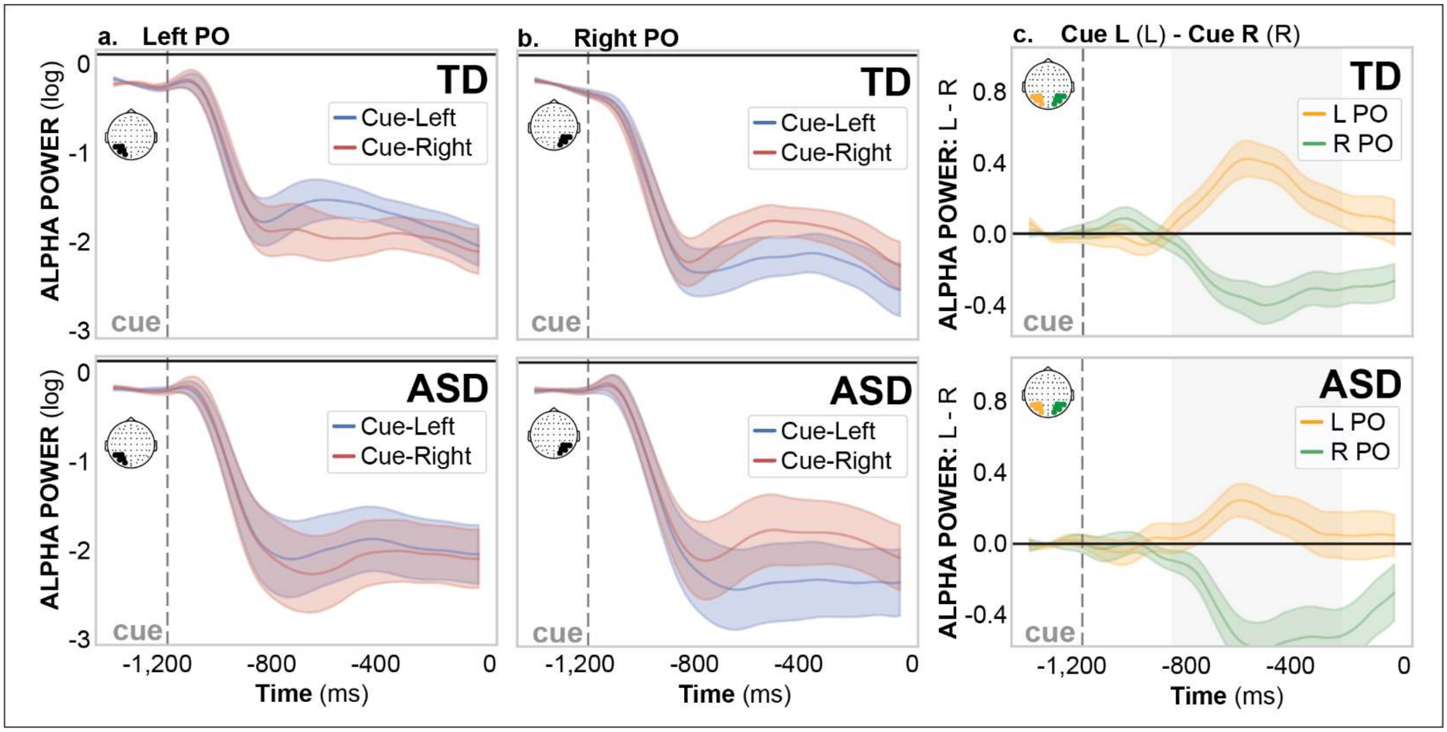


**Supplementary Figure 3. Hemispheric comparison of attentional modulation of alpha-power.**

**a-b.** Time courses of log-transformed alpha-band (7-13 Hz; posterior electrodes) power aligned to cue onset over left parieto-occipital (PO) ROI **(a)** and right PO ROI **(b)**, separated by group (TD: top; ASD: bottom). Cue-left (blue) and cue-right (red) traces are overlaid, with shaded regions representing ±SEM. Thus, within each panel, lines reflect within-hemisphere comparisons of cue direction (i.e., how alpha power differs when attention is directed left vs. right).The time course represents the post-cue window prior to S2, in which the dashed vertical line marks cue onset. The preparatory region, indexing preparation for stimuli, is highlighted by the shaded gray time window (-800 to -200 ms).

**c.** Hemispheric modulation of alpha power are shown as the difference wave between cue-left and cue-right trials (L-R) for both left PO ROI (yellow) and right PO (green), separated by group (TD: top; ASD: bottom). Positive values indicate relatively greater alpha power during leftward compared to rightward orienting, whereas negative values indicate the opposite.

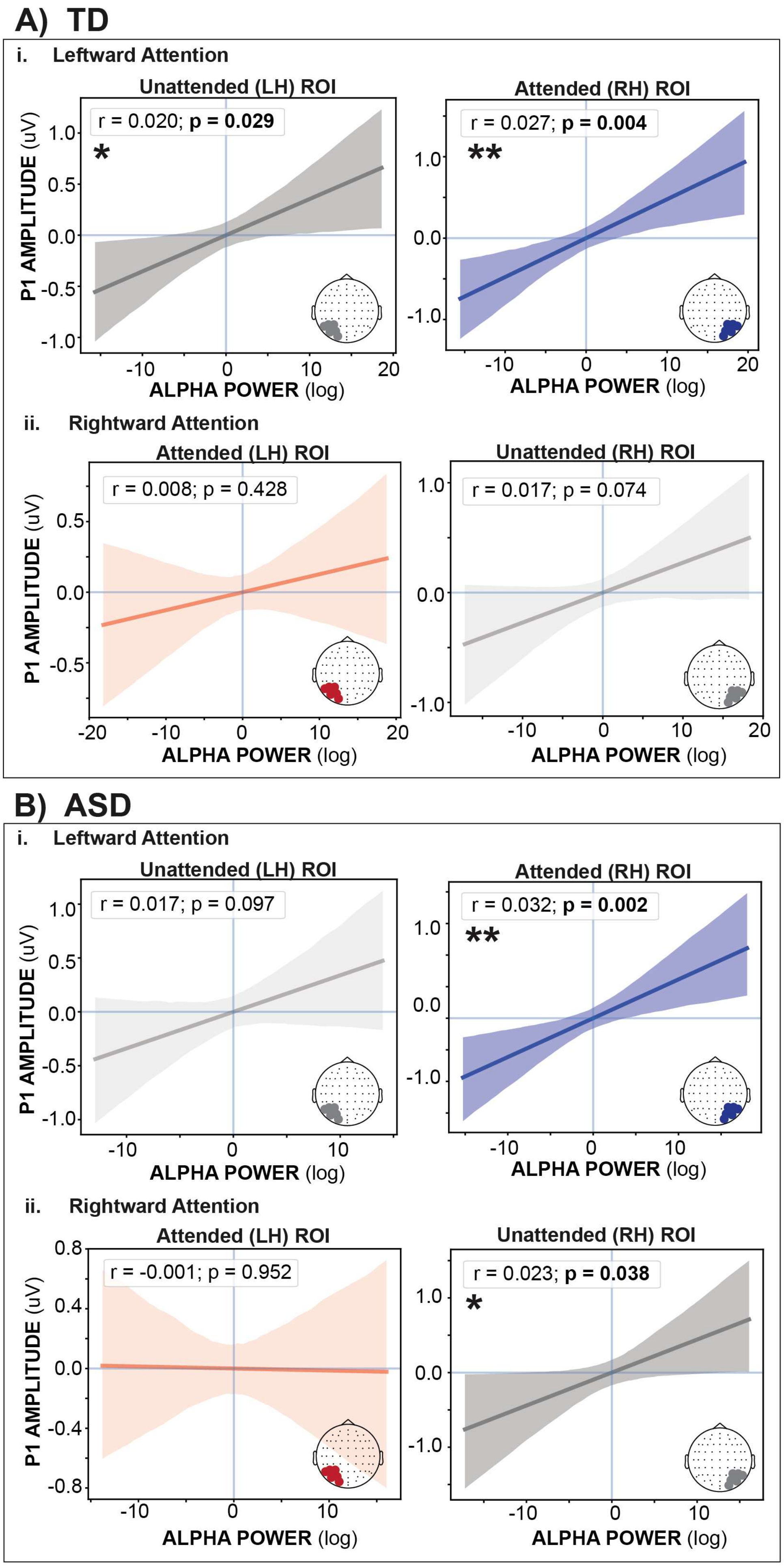


**Supplementary Figure 4. Relationship between anticipatory alpha power and early sensory gain in response to stimulus.**

Correlations between preparatory alpha power and P1 amplitude for TD **(A)** and ASD **(B)** over attended (cue-left: right hemisphere; cue-right: left hemisphere) and unattended (cue-left: left hemisphere; cue-right: right hemisphere) ROI, shown separately for leftward **(i)** and rightward **(ii)** attention. Preparatory alpha power was computed from parieto-occipital electrodes during the −800 to −200 ms pre-stimulus interval, indexing anticipatory sensory gating, and P1 amplitude was extracted from the 110–150 ms post-stimulus window, reflecting early sensory gain of the evoked response. Only trials containing attended stimuli were included to ensure that P1 measurements corresponded to attended processing.

#### Supplementary Tables

| | Group Effects | Cue Side Effects | Stimulus-Type Effects | Block Effects | Age Effects |
|---|---|---|---|---|---|
| **RT** | *None observed* | *None observed* | *None observed* | *None observed* | *None observed* |
| **Hits** | *None observed* | *None observed* | Main effect of Stimulus-Type; driven by lower hit rate to bilateral (vs. unilateral) stimuli | Main effect of Block; driven by lower hit in the inter-mixed (vs. non-social) condition | *None observed* |
| **IES** | *None observed* | *None observed* | *None observed* | Main effect of Block; driven by lower IES in the non-social (vs. social) condition | *None observed* |
| **False Alarms** | *None observed* | *None observed* | Main effect of Stimulus-Type; driven by higher false alarm rate to bilateral (vs. unilateral) stimuli | *None observed* | Main effect of Age; driven by lower false alarm rate in older subjects |
| **d'** | *None observed* | *None observed* | *None observed* | *None observed* | Main effect of Age; driven by higher detection sensitivity in older subjects |

**Supplementary Table 1: Behavioral performance.** Summary of LMM results for behavioral performance, including measures of RT (RT), hit rate, inverse efficiency score (IES), false alarm rate, and detection sensitivity (d’). For each participant, mean RT (RT) was calculated from correct target trials (hits only). Proportion of hits and misses were defined as the presence or absence of a response on target trials, respectively. False-alarm rate was computed as the proportion of non-target trials with a response. Inverse efficiency score (IES) was calculated as mean RT divided by hit rate. Sensitivity (d′) was computed using log-linear–corrected hit and false-alarm rates to avoid ceiling and floor effects. All measures were computed at the participant level. The LMM included fixed effects of Group (TD/ASD) x S2 Type (unilateral/bilateral) × Block (S/NS/IM) × Cue Side (L/R), with Age entered as a covariate, and a random intercept for Subject *(1 | Subject)*.

| | Group Effects | Cue Side Effects | Cue Side x Attention x Social Context Effects | Attention Effects | Social Context Effects | Age Effects |
|---|---|---|---|---|---|---|
| **Pupil** | *None observed* | Main effect of Cue Side; driven by larger pupil response when attention is directed to the right (vs. left) | Cue Side x Attention x Social Context; driven by larger pupil response when attention is directed to the right (vs. left), particularly for unattended social and attended non-social stimuli | *None observed* | *None observed* | *None observed* |

**Supplementary Table 2: Pupil response to unilateral stimuli.** Summary of LMM results for S2-evoked pupil response for non-target unilateral stimuli. Analyses were restricted to non-target trials to eliminate motor-response–related confounds. The LMM included fixed effects of Group (TD/ASD) x Attention (attended/unattended) × Condition (S/NS) × Cue Side (L/R), with Age entered as a covariate, and a random intercept for Subject *(1 | Subject)*.

| | | Attention | | Social Context Effects | Age Effects |
|---|---|---|---|---|---|
| | | **Attention Effects** | **Attention x Cue-Side x Group Interaction** | | |
| **P1** | Amplitude | Main effect of Attention; driven by greater (positive-going) amplitude to attended stimuli | Attention x Cue-Side x Group; driven by greater (positive-going) amplitude to attended stimuli bilaterally in TD, and only for leftward attention in ASD (absent in right) | Main effect of Condition; driven by lower (positive-going) amplitude response to social stimuli | Main effect of Age; driven by lower (positive-going) amplitude in older subjects |
| | Latency | *None observed* | Attention x Cue-Side x Group; driven by earlier latency to left (vs. right) visual-field stimuli in TD only, with the ASD group exhibiting later latencies to left visual-field stimuli (vs. TD) | Main effect of Condition; driven by earlier latency in response to social stimuli | Main effect of Age; driven by earlier latency responses in older subjects |

**Supplementary Table 3: Evoked response to unilateral stimuli.** Summary of LMM results for S2-evoked P1 component for non-target unilateral stimuli. Analyses were restricted to non-target trials to eliminate motor-response–related confounds. The LMM included fixed effects of Group (TD/ASD) x Attention (attended/unattended) × Condition (S/NS) × Cue Side (L/R), with Age entered as a covariate, and a random intercept for Subject *(1 | Subject)*.

| | Attention | | | | |
|---|---|---|---|---|---|
| | **Attention Effects** | **Attention: Interaction between Attention and Cue Side** | **Attention: Interaction between Attention and Cue Side and Group** | **Block Effects** | **Age Effects** |
| **Alpha power** | Main effect of Attention; driven by greater alpha power over unattended ROI (vs. attended ROI) | Attention x Cue Side; driven by greater alpha power over unattended ROI (vs. attended ROI), only when attention is directed to the left | Attention x Cue Side x Group; both groups showed robust attentional modulation during leftward attention, but only ASD exhibited greater alpha power over unattended ROI (vs. attended during rightward attention; TD exhibited bilateral cue-side lateralization, with lower alpha power over the attended ROI and higher alpha power over the unattended ROI during leftward (vs. rightward) attention, whereas ASD showed cue-side modulation restricted to the attended ROI, with no significant modulation of the unattended ROI. | Main effect of Block; driven by lower alpha power in the inter-mixed blocks | *None observed* |

**Supplementary Table 4: Anticipatory alpha band neuro-oscillatory activity.** Summary of LMM model results for anticipatory alpha power (-800 to -200 ms) for all trials. The LMM included fixed effects of Group (TD/ASD) x Attention (unattended/attended ROI) × Block (S/NS/IM) × Cue Side (L/R), with Age entered as a covariate, and a random intercept for Subject *(1 | Subject)*.
